# Zinc-finger protein 518 plays a crucial role in pericentromeric heterochromatin formation by linking satellite DNA to heterochromatin

**DOI:** 10.1101/2022.09.15.508097

**Authors:** Shinya Ohta, Jun-Ichirou Ohzeki, Nobuko Sato, Ken-Ichi Noma, Hiroshi Masumoto

## Abstract

Aneuploidy is caused by chromosomal missegregation and is frequently observed in cancers and hematological diseases. Pericentromeric heterochromatin formation is essential for proper chromosome segregation; however, it is unclear how heterochromatin is targeted to pericentromeres. In this study, we investigated the involvement of two homologous zinc-finger proteins (ZNF518A and ZNF518B) in heterochromatin formation at pericentromeres by determining the cellular localization of ZNF518, centromeric proteins such as CENP-A and CENP-B, and heterochromatin factors, including HP1 and histone H3K9 trimethylation. The results showed that various segments in ZNF518 interact with CENP-B, HP1, and G9a histone H3K9 methyltransferase. In conclusion, we identified a novel mechanism underlying pericentromeric heterochromatin formation mediated by ZNF518.

## Introduction

Defects in centromere or kinetochore functions are related to chromosomal abnormalities, and anomalous chromosome segregation can lead to cancer or hematological diseases.^1,2^ Chromosomal abnormalities can occur during mitosis, where replicated chromosomes are normally equally distributed between two daughter cells. During mitosis, compacted chromosomes are segregated when the spindle attaches to and pulls the kinetochore, a proteinaceous structure on the centromere. Centromere protein (CENP)-A, a centromere-specific histone H3 variant, is essential for kinetochore formation and directs the assembly of a complex containing at least 15 other proteins, including CENP-C.^3^ CENP-A, CENP-B, and CENP-C are evolutionarily conserved from fission yeast to vertebrates and show distinct association patterns within the centromere.^4, 5^

In normal human chromosomes, centromeres assemble on α-satellite DNA (alphoid DNA), a giant repetitive DNA locus consisting of 171-bp repeat units.^6, 7^ CENP-A assembles on a portion of α-satellite DNA, forming an epigenetic mark of kinetochore assembly.^8–11^ Some α-satellite DNA subclasses contain a 17-nucleotide CENP-B box, bound by CENP-B.^8, 12, 13^ Thus, CENP-B is distributed in both CENP-A chromatin (core centromere) and pericentromeric heterochromatin. CENP-B is involved in the assembly of CENP-A and CENP-C in centromeres and is crucial for functional centromere formation.^9, 10^ Furthermore, CENP-B and CENP-B boxes are required for human artificial chromosome formation and *de novo* centromeric CENP-A chromatin assembly in cells with synthetic alphoid DNA.^14–16^ CENP-B also induces heterochromatin formation in alphoid DNA integrated into chromosomal arms.^16^ Interestingly, CENP-B recruits numerous factors, including the open chromatin modifier ASH1L (H3K36 methylase) and heterochromatin factor HP1, in a mutually exclusive manner.^17^ Heterochromatin plays important roles in chromosome stability and the maintenance of sister chromatid cohesion during chromosome segregation.^18, 19^ Therefore, both centromeric chromatin and heterochromatin are important for accurate chromosome segregation.

Pericentromeric heterochromatin insulates centromeres from the rest of the genome and is required for proper mitotic chromosome segregation.^19, 20^ Heterochromatin has a more condensed structure than euchromatin and carries specific post-translational histone modifications. Nucleosomal surfaces with charge differences contribute to the condensed structure of heterochromatin.^21–23^ Histone H3 lysine 9 trimethylation (H3K9me3) is recognized and bound by the chromodomain of the heterochromatin protein HP1.^24^ HP1 in turn recruits the H3K9 methyltransferase, suppressor of variegation 3-9 homolog (SUV39H).^18, 25^ The HP1-mediated loading of SUV39H to chromatin is required for the maintenance of heterochromatin at pericentromeres.^26, 27^ As SUV39H localizes to pericentromeres in addition to other constitutive heterochromatin regions, heterochromatin formation at pericentromeres involves SUV39H. The mechanisms by which heterochromatin factors are initially targeted to pericentromeres, however, remain elusive.^28^

H3K9 methyltransferases other than SUV39H have been identified. SET domain bifurcated 1 (SETDB1) catalyzes H3K9me3 formation, whereas euchromatic histone lysine methyltransferases 1 and 2 (GLP and G9a) are responsible for H3K9me1 and H3K9me2, respectively. G9a can also methylate Lys9 and Lys27 of histone H3.^29^ These H3K9 methyltransferases, including SUV39H, are selectively loaded onto distinct genomic regions through protein interactions with other factors.^30^ G9a and GLP bind to the widely interspaced zinc-finger (WIZ) protein.^29, 31, 32^ H3K9 methylation at chromosomal arm regions is mostly lost in G9a-deficient cells, whereas H3K9 methylation at pericentromeres is largely lost in SUV39H-deficient cells.^29^ Moreover, G9a interacts with zinc-finger protein 518B (ZNF518B) as recently indicated by a proteomics analysis.^32^ Our previous proteomics analysis revealed a possible role for ZNF518B as a functional protein on mitotic chromosomes.^33, 34^ The present study demonstrated that G9a, which acts as a H3K9 methyltransferase for chromosomal arm regions, is also involved in pericentromeric heterochromatin formation via interaction with ZNF518B recruited by CENP-B.

## Results

### Human ZNF518A/B is recruited to centromeres by CENP-B

We first examined the localization of green fluorescent protein (GFP)-tagged ZNF518B and CENP-A, CENP-B, and CENP-C in human U2OS cells. Overexpression of GFP fusion proteins allows for visualization of even weakly expressed proteins; using this technique, we were able to successfully visualize the localization of ZNF518. GFP-ZNF518B co-localized proximally to the centromeric factors (Fig. 1A–F). ZNF518B localized to the centromeres during mitosis (Fig. S1A), suggesting constitutive centromeric localization of ZNF518B throughout the cell cycle. GFP-ZNF518B foci were slightly shifted away from CENP-A and CENP-C foci, but coincided with CENP-B foci, indicating that ZNF518B localizes to the pericentromeric region, where it is bound by CENP-B (Fig. 1A–F).

**Figure 1.**
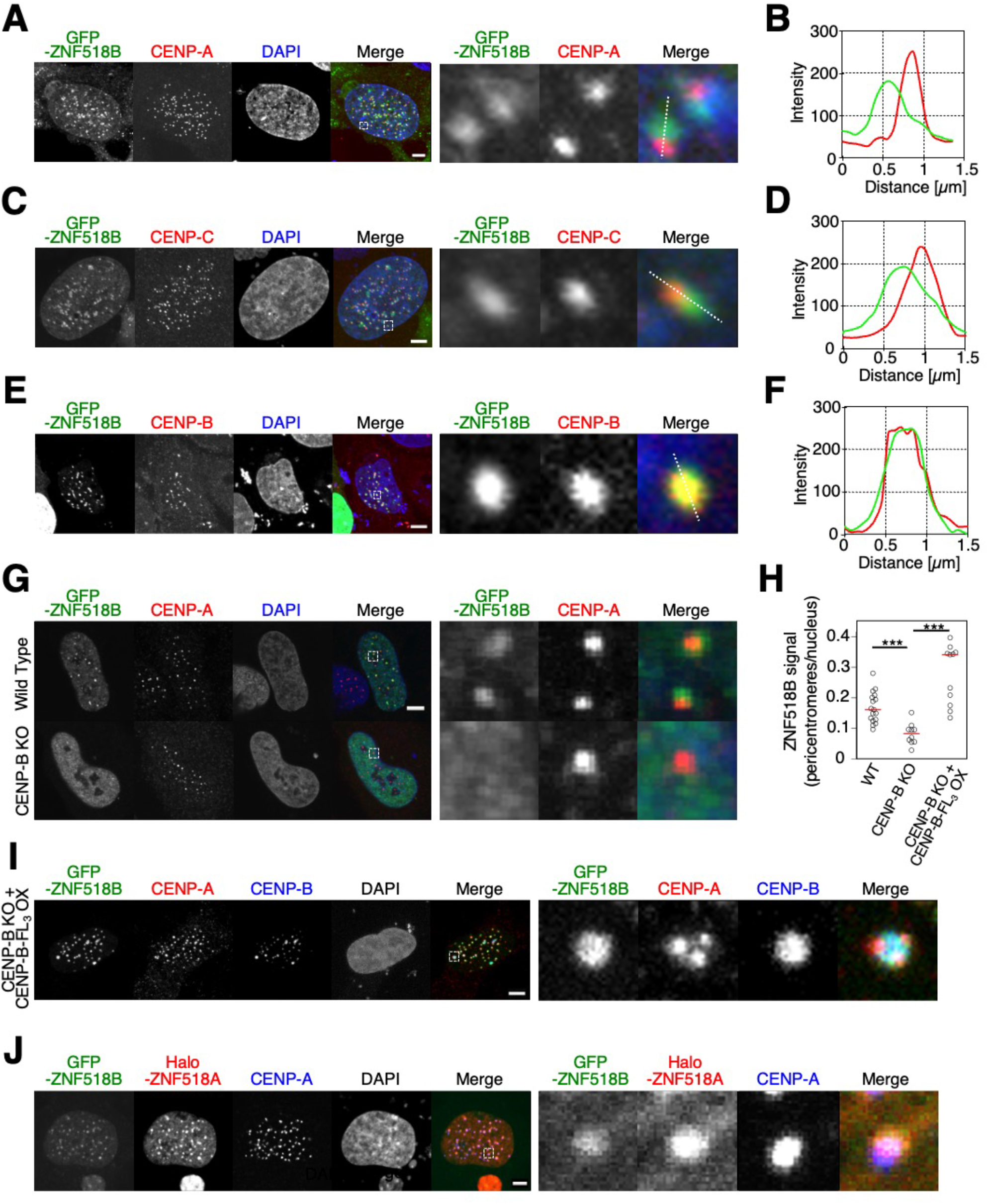
ZNF518B and ZNF518A localize to centromeres in a CENP-B-dependent manner. **(A, C, and E)** Co-localization of GFP-ZNF518B (green) and the centromeric factors (red) CENP-A **(A),** CENP-B **(C),** and CENP-C **(E)**. Areas marked by dashed boxes in the left images are magnified in the right panel. Scale bars, 5 μm. **(B, D, and F)** Line plot analyses along the white lines in panels A, C, and E for quantification of GFP-ZNF518B (green) and the centromeric factors (red) CENP-A **(B)** CENP-B **(D),** and CENP-C **(F)**. **(G)** Localization of GFP-ZNF518B (green) and CENP-A (red) in WT (upper) and *CENPB*-KO cells (lower). Areas marked by dashed boxes in the left panels are magnified in the right panels. **(H)** Quantification of the fluorescence intensities of GFP-ZNF518B signals at centromeres in WT, *CENPB*-KO [see panel **(G)**], and *CENPB*-KO cells overexpressing CENP-B-Flag_3_ [*CENPB*-KO + CENP-B-FL_3_ OX; see panel **(I)**]. The y-axis shows the relative fluorescence intensity of GFP-ZNF518B signals associated with centromeres to total GFP signals in the nucleus. Circles represent measurements from the individual cells shown in panels G and I. Red bars indicate medians. \*\*\**p* < 0.005. **(I)** Co-localization of GFP-ZNF518B (green), CENP-A (red), and CENP-B (blue) in *CENPB*-KO cells overexpressing CENP-B-Flag_3_ (*CENPB*-KO + CENP-B-FL_3_ OX). The area marked by the dashed box in the left panel is magnified in the right panel. **(J)** Co-localization of Halo-ZNF518A (red), GFP-ZNF518B (green), and CENP-A (blue). The area marked by the dashed box in the left panel is magnified in the right panel.

To investigate the localization dependency between ZNF518B and CENP-B, we examined the localization of GFP-ZNF518B in *CENPB*-knockout (KO) HeLa cells.^17^ Centromeric/pericentromeric localization of GFP-ZNF518B was lost in *CENPB*-KO cells, but was restored upon exogeneous CENP-B expression (Fig. 1G– I), indicating that the centromeric localization of ZNF518B depends on CENP-B.

ZNF518A is a paralogue of ZNF518B present in humans (Fig. S1B). To determine the localization of ZNF518A, GFP-ZNF518B and Halo-ZNF518A were transiently co-expressed in U2OS cells (Fig. 1J). GFP-ZNF518B and Halo-ZNF518A foci co-localized at the centromeres. Again, the co-localized signals were slightly shifted away from CENP-A foci, suggesting that both ZNF518A and ZNF518B localize at centromeres (Fig. 1J). Centromeric localization of GFP-ZNF518A was lost in *CENPB*-KO cells, but restored upon exogenous CENP-B expression (Fig. S1C), indicating that the centromeric localization of ZNF518A depends on CENP-B. Taken together, the results suggested that both ZNF518A and ZNF518B localize to the pericentromere via a common molecular mechanism involving CENP-B (Fig. S1D).

### The C-terminus of ZNF518A/B is required for centromeric localization

To unravel the molecular mechanisms underlying ZNF518 recruitment to centromeres, we sought to identify the segments in the 1,074-amino-acid (aa) ZNF518B protein that are required for centromeric localization. We examined the centromeric localization of 15 ZNF518B derivatives expressed as GFP fusion proteins (Fig. 2). ZNF518B derivatives containing regions from the N-terminus to 628 aa (ZNF518B^1–398^, ZNF518B^344–628^, and ZNF518B^1–628^) did not show centromeric localization (Fig. 2, S2A). Therefore, the N-terminus of ZNF518B does not form sufficient interactions to localize the protein to centromeres. Among four ZNF518B derivatives with incrementing C-terminal truncations (ZNF518B^344–1074^, ZNF518B^344–987^, ZNF518B^344–877^, and ZNF518B^344–769^), ZNF518B^344–1074^, ZNF518B^344–987^, and ZNF518B^344–877^ showed centromeric localization in 70.7%, 58.9%, and 57.1% of transfected cells, respectively, whereas ZNF518B^344–769^ did not show centromeric localization (Fig. 2). Similarly, among ZNF518B^555–1074^, ZNF518B^555–877^, and ZNF518B^555–769^, only ZNF518B^555–769^, which had the same C-terminal truncation as ZNF518B^344–769^, did not show centromeric localization (Figs. 2, S2A). These results suggested that residues 770–877 of ZNF518B are involved in its centromeric localization. However, only 2.2% of transfected cells showed centromeric localization of ZNF518B^770–877^ (Fig. 2). Therefore, residues 770–877 are important, but insufficient for centromeric localization.

**Figure 2.**
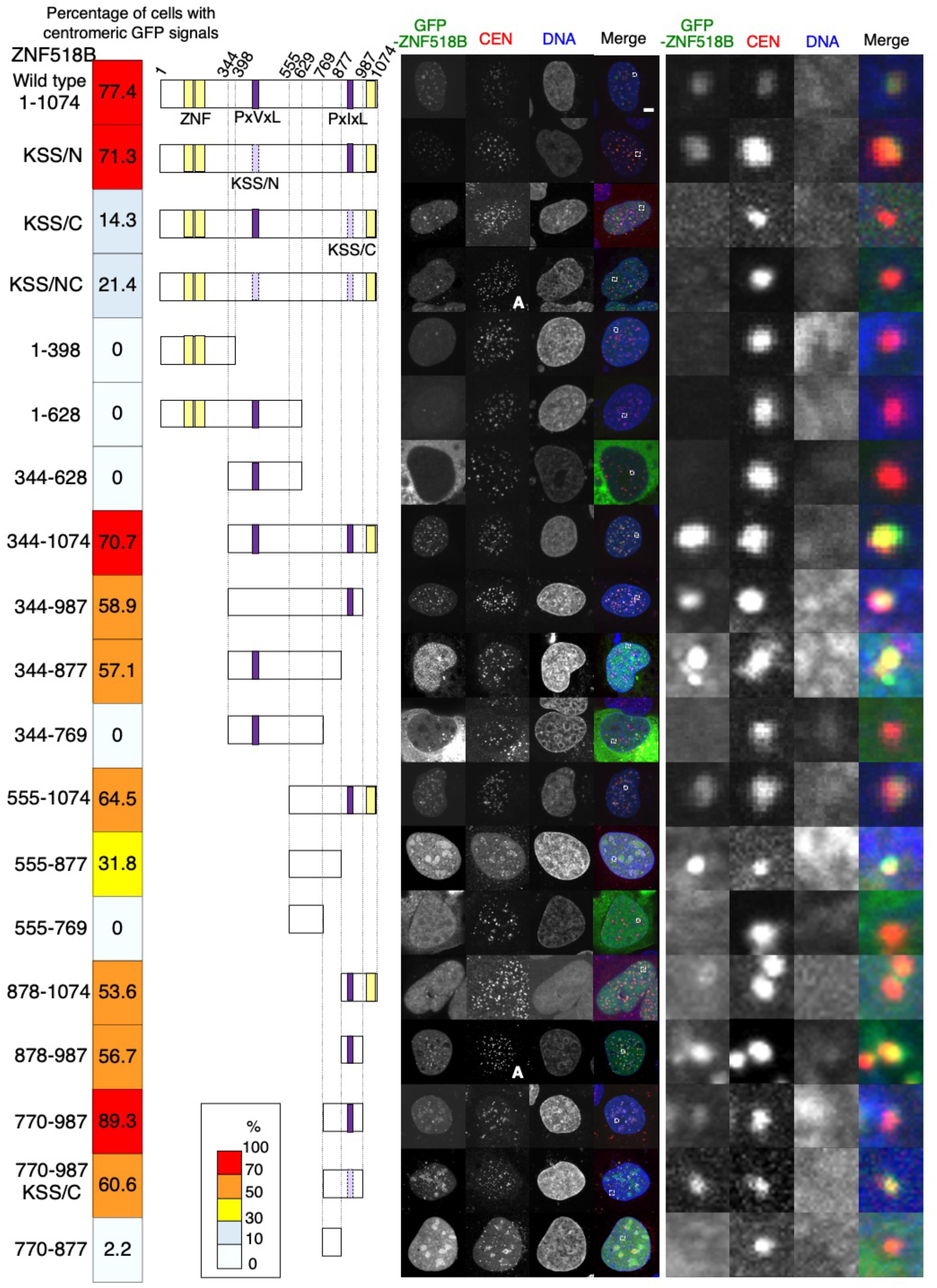
Identification of ZNF518B segments required for centromeric localization. Localization of the ZNF518B derivatives at centromeres. The schematic representation of the respective ZNF518B derivatives is drawn approximately to scale. Yellow boxes indicate zinc-finger motifs. The two PxV(I)xL motifs (purple boxes) were substituted with KxSxS (light purple boxes), shown as KSS/N and KSS/C. KSS/NC indicates that both PxV(I)xLs motifs were substituted with KxSxS. GFP-fused ZNF518B signals (WT and 18 derivatives; green) were co-visualized with the centromeric marker CENP-A or CENP-B (left images). The two images (ZNF518B^KSS/NC^ and ZNF518B^878–987^) labeled with “A” show CENP-A foci, all other images show CENP-B signals. Areas marked by dashed boxes in the left images are magnified in the right panel. Numbers and colors (left) reflect % frequencies of cells with GFP-ZNF518B signals at pericentromeres. Scale bar, 5 μm.

Given the large difference in centromeric localization among ZNF518B^555–1074^, ZNF518B^555–877^, and ZNF518B^555–769^ (in 64.5%, 31.8%, and 0% of transfected cells, respectively), we examined the centromeric localization of ZNF518B derivatives carrying residues 770–1074 (Figs. 2, S2A). ZNF518B^878–1074^, ZNF518B^878–987^, and ZNF518B^770–987^ showed centromeric localization in 53.6%, 56.7%, and 89.3% of cells, respectively (Figs. 2, S2A). ZNF518B^770–987^ showed the highest frequency of centromeric localization among all ZNF518B derivatives. These results suggested that residues 770–987 are involved in the centromeric localization of ZNF518B.

We next examined the centromeric localization of ZNF518A derivatives (Fig. S2B). Wild-type (WT) ZNF518A (ZNF518A^WT^) and two derivatives containing the C-terminal 1317–1483-aa segment (ZNF518A^971–1483^ and ZNF518A^1317–1483^) showed high frequencies of centromeric localization, suggesting that the C-terminal portion of ZNF518A harbors crucial segment(s) for centromeric localization (Fig. S2B). Notably, the C-terminal segments of ZNF518A and ZNF518B required for centromeric localization carry PxV(I)xL HP1-interaction motifs, which are widely conserved among higher vertebrates, including fish (Fig. S3), implying that their role is evolutionarily conserved potentially through their localization at centromeres.

### The C-terminal PxIxL motif is required for the centromeric localization of ZNF518B

To elucidate the functional significance of the PxV(I)xL motifs in ZNF518B, we constructed the GFP-ZNF518B^KSS/C^ derivative in which the conserved hydrophobic residues PxIxL in the C-terminus were substituted with the hydrophilic residues KxSxS (Fig. 2). ZNF518B^KSS/C^ exhibited less frequent centromeric localization (14.3%) than ZNF518^WT^ (77.5%; Fig. 2). In contrast, the centromeric localization of ZNF518B^KSS/N^, which carried a substitution of the PxVxL motif with KxSxS in the N-terminus, was comparable to that of ZNF518^WT^ (Fig. 2). ZNF518B^KSS/NC^, which carried KxSxS substitutions of the PxV(I)xL motifs in both the N- and C-termini, showed a similar centromeric localization frequency (21.4%) as ZNF518B^KSS/C^ (Fig. 2). Therefore, the PxIxL motif present in the C-terminus is involved in the centromeric localization of ZNF518B. ZNF518A^KSS/C^ also showed reduced centromeric localization (Fig. S2B). These results demonstrated that the conserved PxV(I)xL motif in the C-terminus of ZNF518 is important for centromeric localization.

Although the C-terminal PxIxL motif is important for centromeric localization, ZNF518B^KSS/C^ and ZNF518B^KSS/NC^ localized to the centromeres in 14.3% and 21.4% of cells, respectively, implying the involvement of other interactions in centromeric localization. To test this, we substituted the PxVxL motif with KxSxS in the ZNF518B^770–987^ derivative that localized to centromeres with the highest frequency (89.3%), yielding ZNF518B^770–987^ ^KSS/C^. ZNF518B^770–987^ ^KSS/C^ showed centromeric localization in 60.6% of transfected cells (Fig. 2, S2B), indicating that both the PxIxL motif and the C-terminal segment (770–987 aa) participate in the centromeric localization of ZNF518B.

### ZNF518B interacts with HP1α and HP1β

The localization of full-length ZNF518B^WT^ to centromeres and pericentromeres requires CENP-B. The ZNF518B^770–987^ derivative with the C-terminal segment (770– 987 aa) including the PxIxL motif also localized to centromeres (Fig. 2). In *CENPB*-KO cells, ZNF518B^770–987^ showed a focal localization that was not observed for ZNF518B^WT^ (Fig. S4A). However, the ZNF518B^770–987^ foci were distant from centromeres and rather co-localized with heterochromatin foci in *CENPB*-KO cells (Fig. S4B). Thus, ZNF518B^770–987^ can be recruited to non-centromeric heterochromatin domains through its interaction with heterochromatic factors in the absence of CENP-B (Fig. S4C).

Given its similarity to the canonical HP1-interacting motif PxVxL, we hypothesized that the PxIxL motif in the C-terminus of ZNF518B interacts with HP1, resulting in the accumulation of ZNF518B at heterochromatin regions in *CENPB*-KO cells. Three isozymes of human HP1 have been identified: HP1α, HP1β, and HP1γ.^35^ We evaluated the interactions between ZNF518B and the HP1 isoforms using the fluorescence microscopy-based interaction-trap (FMIT) assay,^36^ which can be used to examine protein interactions on chromatin *in vivo*. Given the overabundance of target proteins compared to their endogenous counterparts, this technique allows for the detection of direct and indirect protein–protein interactions using fluorescence microscopy.^17, 20, 36–40^

The FMIT assay was performed using HeLa-Int-03 cells, in which approximately 15,000 repeats of a synthetic alphoid DNA dimer unit (derived from the human chromosome 21 centromere) containing the *tet*O sequence (alphoid*^tetO^*) were ectopically integrated into a chromosome arm region, and *CENPB* was knocked out (Fig. 3A).^17^ TetR-EYFP-ZNF518B localized to the alphoid*^tetO^* repeats, but did not exhibit centromeric localization due to the absence of CENP-B. We next examined whether Halo-HP1 co-expressed with TetR-EYFP-ZNF518B would be recruited to the alphoid*^tetO^* array (Fig. 3B). Halo-HP1α, -HP1β, and -HP1γ were recruited to alphoid*^tetO^* in 48%, 58%, and 14% of cells, respectively (Fig. 3C). In negative control cells expressing TetR-EYFP not fused to ZNF518B, Halo-HP1α, -HP1β, and -HP1γ were barely recruited to alphoid*^tetO^* (Fig. 3C). These results suggested that ZNF518B interacts with HP1.

**Figure 3.**
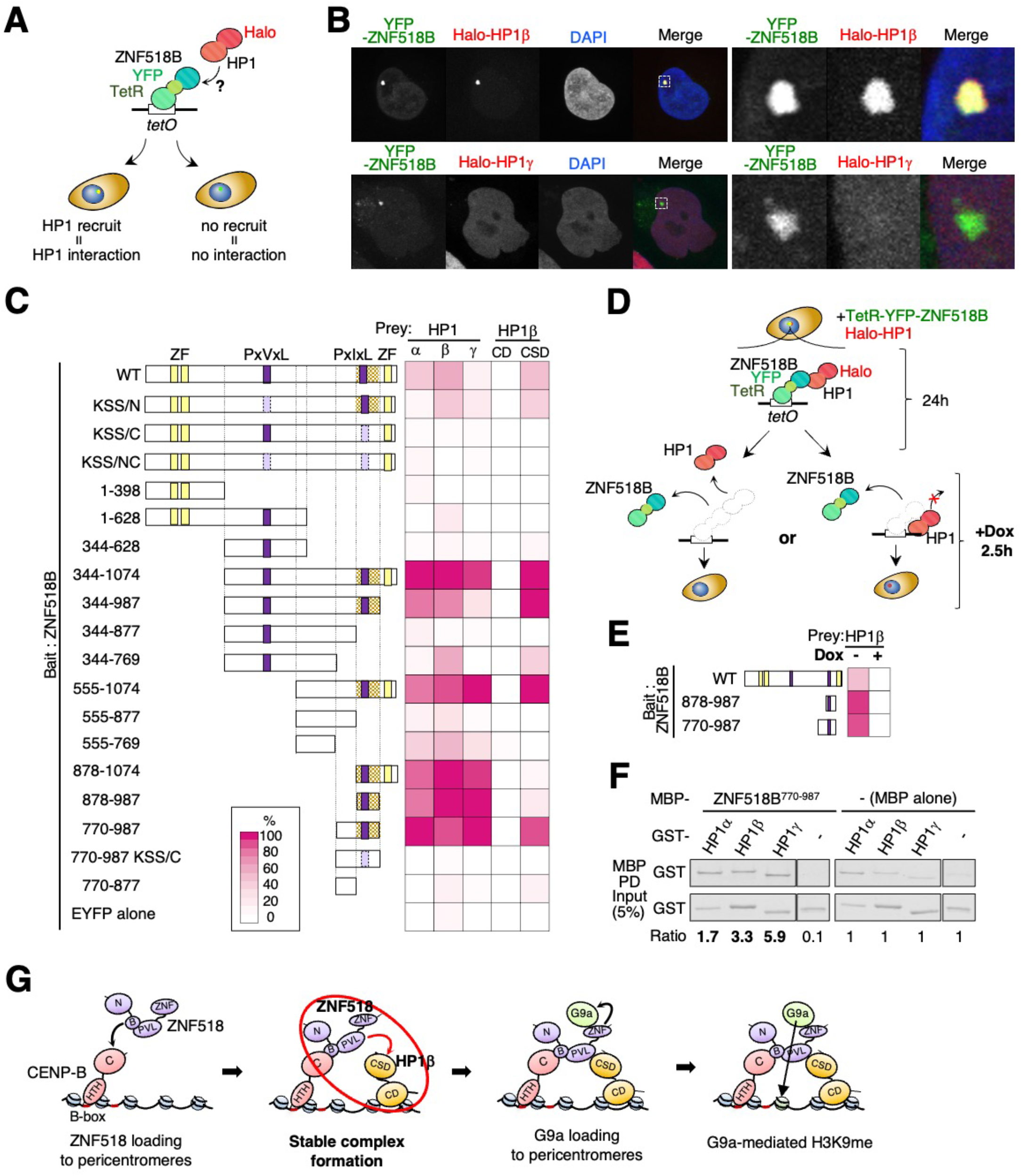
The PxIxL motif in ZNF518B is required for interaction with the chromo-shadow domain of HP1. **(A)** Schematic diagram of the FMIT assay for investigating the interaction between ZNF518B and HP1. *CENPB*-KO cells were used. **(B)** Tethering of YFP-ZNF518B (green) to the *tetO* repeats results in the accumulation of Halo-HP1β (top; red), but not Halo-HP1γ (bottom; red) at the locus. **(C)** FMIT assay to examine the interaction between ZNF518B derivatives and HP1. Cells expressing TetR-EYFP-ZNF518B (WT or derivatives) and Halo-HP1 were used. The ZNF518B segment predicted to interact with HP1 is indicated by shaded boxes. Numbers and colors reflect % frequencies of the co-localization of TetR-EYFP-ZNF518B with full-length HP1 (HP1α, HP1β, and HP1γ) and the chromo domain (CD) and chromo-shadow domain (CSD) of HP1β. **(D)** Experimental design to examine whether ZNF518B is required to maintain HP1β localization at the *tetO* repeats. TetR-EYFP-ZNF518 quickly dissociated from the *tetO* repeats in the presence of Dox. *CENPB* KO cells were used in this assay. **(E)** Cells carrying TetR-EYFP-ZNF518 (WT or derivatives) and Halo-HP1β were subjected to the FMIT assay as explained in panel D. **(F)** Pull-down (PD) experiments to examine the physical interaction between ZNF518B and HP1. Purified recombinant GST-HP1 (input) was incubated with MBP-ZNF518B^770–987^, and MBP PD samples were subjected to immunoblotting using anti-GST. Signal intensities in the indicated samples were normalized to those in samples with MBP alone. **(G)** Model depicting the central roles of ZNF518A and ZNF518B in pericentromeric heterochromatin formation via their interactions with CENP-B, HP1, and G9a. The red ellipse and arrow highlight the reaction step focused on in this figure.

### ZNF518B interacts with HP1 isoforms via the C-terminal PxIxL motif

We next investigated which segments in ZNF518B are required for interaction with the HP1 isoforms by subjecting the ZNF518B derivatives described in Figure 2 to the FMIT assay (Fig. 3C). Upon co-expression with TetR-EYFP-fused ZNF518B derivatives (ZNF518B^1–398^, ZNF518B^1–628^, ZNF518B^344–628^, ZNF518B^344–877^, ZNF518B^555–877^, and ZNF518B^770–877^), Halo-HP1 was scarcely or not recruited to alphoid*^tetO^*. As these derivatives lacked the PxIxL motif, this result indicated the importance of the PxIxL motif in the interactions between ZNF518B and the HP1 isoforms (Fig. 3C).

We further examined the significance of the PxV(I)xL motifs in the ZNF518B-HP1 interactions using TetR-EYFP-fused ZNF518B derivatives (ZNF518B^KSS/N^, ZNF518B^KSS/C^, ZNF518B^KSS/NC^, and ZNF518B^770–987^ ^KSS/C^) carrying PxV(I)xL substitutions with KxSxS, as depicted in Figure 2. Halo-HP1 was scarcely recruited to alphoid*^tetO^* by the ZNF518B derivatives, except for ZNF518B^KSS/N^. Therefore, the PxIxL motif in the C-terminus, not the PxVxL motif in the N-terminus, participates in the interaction with HP1 (Fig. 3C).

Next, the TetR-EYFP-fused ZNF518B derivatives ZNF518B^344–1074^, ZNF518B^344–987^, ZNF518B^555–1074^, ZNF518B^878–1074^, ZNF518B^878–987^, and ZNF518B^770–987^, carrying the PxIxL motif in the ZNF518B C-terminus, were subjected to the FMIT assay. Halo-HP1 was recruited to alphoid*^tetO^* in more than 75% of cells (Fig. 3C). Remarkably, these N-terminal-deleted derivatives were able to recruit the three HP1 proteins with similar frequencies. Furthermore, ZNF518B^344–1074^ was more efficient at recruiting HP1 than ZNF518B^WT^. Therefore, the N-terminal segment (1– 398) is inhibitory for ZNF518B interactions with HP1 isoforms (Fig. 3C).

### The ZNF518B PxIxL motif interacts with the chromo-shadow domain of HP1

HP1 comprises a central linker flanked by chromo and chromo-shadow domains, the latter of which interacts with the PxVxL motif in CAF1/p150.^41^ As the PxIxL motif in ZNF518B is crucial for interaction with HP1 isoforms, we predicted that ZNF518B interacts with the chromo-shadow domain rather than the chromo domain of the HP1 isoforms. We examined the interactions between ZNF518B and the chromo (HP1β^1– 92^) and chromo-shadow (HP1β^93–185^) domains of HP1β using the FMIT assay. As expected, only the chromo-shadow domain was recruited to alphoid*^tetO^* by ZNF518B derivatives carrying the PxIxL motif in the C-terminus. This confirmed the interaction between the PxIxL motif of ZNF518B and the chromo-shadow domain of HP1β (Fig. 3C).

ZNF518B^344–769^ and ZNF518B^555–769^ weakly bound HP1β despite the absence of a PxIxL motif. The ZNF518B PxIxL motif had similar binding affinities for all three HP1 isoforms. However, ZNF518A residues 971–1321, which correspond to ZNF518B residues 555–769, did not show such specificity for the HP1 isoforms (Fig. S5B). Therefore, the 555–769-aa segment in ZNF518B is a subsidiary segment for HP1 interactions that determine its specificity for HP1β.

### ZNF518B maintains HP1 at alphoid*^tetO^*

We next examined whether HP1β recruited to alphoid*^tetO^* by TetR-EYFP-ZNF518B is maintained at the alphoid*^tetO^* repeat locus when ZNF518B is released from the alphoid*^tetO^* DNA following the addition of doxycycline (Dox) (Fig. 3D). TetR-EYFP-ZNF518 was rapidly released from alphoid*^tetO^*after the addition of Dox. EYFP foci disappeared in 96% of cells 120 min after Dox addition (Fig. S5C). We quantified Halo-HP1β foci at alphoid*^tetO^* 150 min after Dox addition (Fig. 3E). The Halo-HP1β foci disappeared from the alphoid*^tetO^* DNA in all assays employing ZNF518B^WT^, ZNF518B^878–987^, and ZNF518B^770–987^ (Fig. 3E). Therefore, ZNF518B is required to maintain HP1β at the alphoid*^tetO^*.

### ZNF518B interacts directly with HP1 isoforms *in vitro*

The FMIT results strongly suggested that ZNF518B interacts with the HP1 isoforms, corroborating previously published results regarding the interaction of ZNF518B with the HP1 isoforms.^42^ To further examine their direct interactions, we purified the ZNF518B^770–987^ derivative containing the PxIxL motif as a maltose-binding protein (MBP) fusion protein and the HP1 isoforms as glutathione S-transferase (GST) fusion proteins, respectively, using an *Escherichia coli* overexpression system. The two purified proteins were mixed, and the MBP-fused ZNF518B^770–987^ proteins were pulled down using magnetic amylose beads. The GST-HP1 isoforms were co-purified with ZNF518B^770–987^ and were more abundant in pull-down (PD) samples with MBP-ZNF518B^770–987^ than in those with MBP alone (Fig. 3F), suggesting that the PxIxL motif in the ZNF518B C-terminus directly interacts with HP1α, HP1β, and HP1γ (Fig. 3G).

### ZNF518B localization cannot be maintained at centromeres without CENP-B

As CENP-B is required for centromeric localization of ZNF518B (Fig. 1), we next investigated whether it is required to maintain ZNF518B at the centromeres. We created a *CENPB* conditional KO (cKO) system using U2OS cells (Fig. S6A). The *Oryza sativa TIR1* gene, encoding an auxin receptor, was inserted into an adeno-associated virus. In virus-infected cells, endogenous CENP-B fused with a mini auxin-inducible degron (mAID) tag was rapidly targeted for degradation via the ubiquitin-proteasome pathway following indole-3-acetic acid (IAA) treatment (Fig. S6A).^43^

The localization of GFP-ZNF518B was analyzed under three conditions (Fig. 4A): CENP-B:mAID available at all times (CENP-B^ON^); CENP-B:mAID removed 24 h before ZNF518B transfection (CENP-B^OFF^); and CENP-B:mAID removed 4 h before fixation (CENP-B^4h-OFF^). As expected, GFP-ZNF518B localized at the centromeres in CENP-B^ON^ cells (Fig. 4B–E). In CENP-B^OFF^ cells, CENP-B signals at the centromeres were below the detection limit, and GFP-ZNF518B dissociated from the centromeres (Fig. 4B–E). In CENP-B^4h-OFF^ cells, GFP-ZNF518B also dissociated from the centromeres (Fig. 4B–E). These results demonstrated that the recruitment and maintenance of ZNF518B at centromeres require CENP-B.

**Figure 4.**
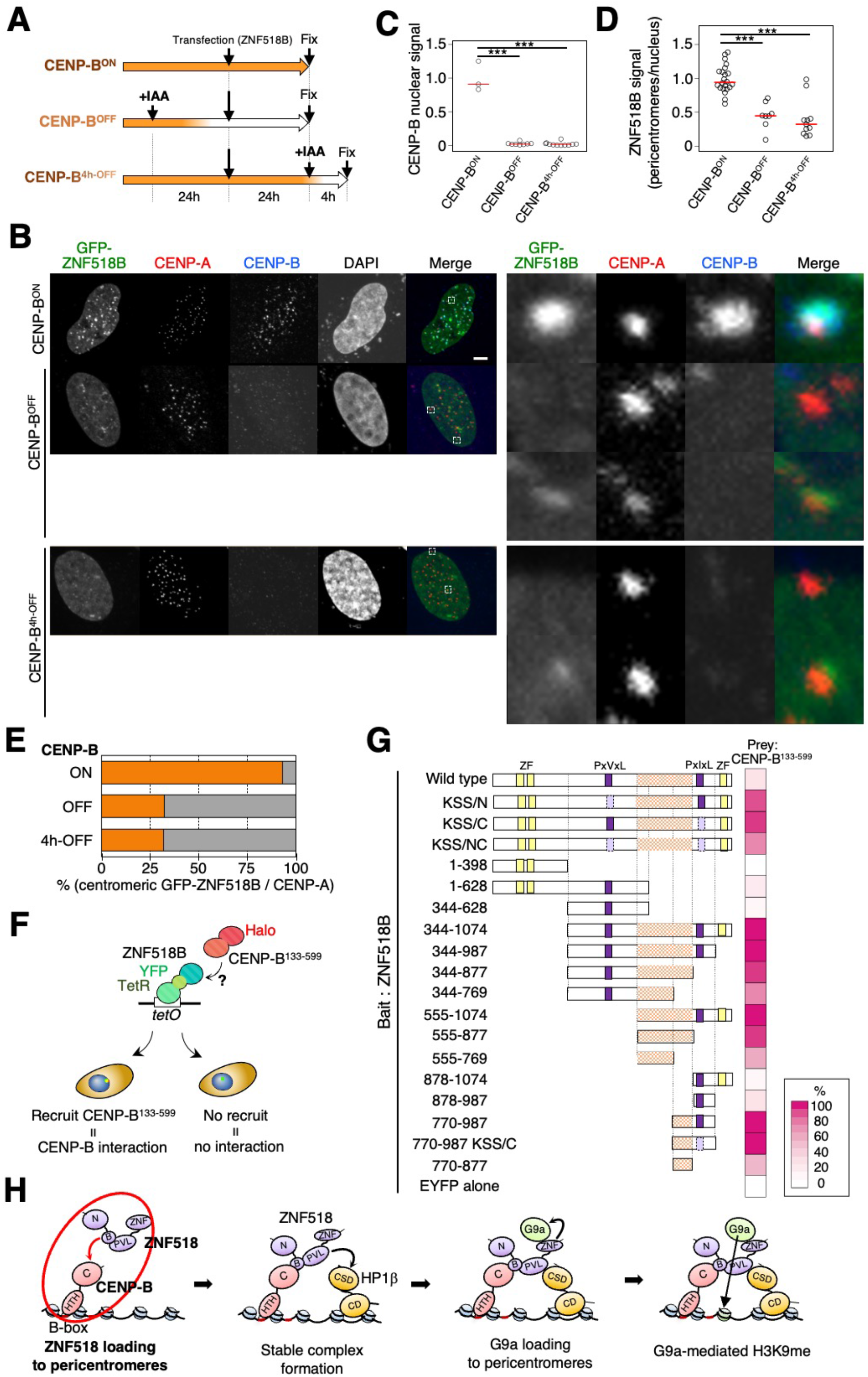
Dissection of the interaction of ZNF518B with CENP-B. **(A)** Schematic representation of the GFP-ZNF518B localization assay, in which indole-3-acetic acid (IAA) was added at different time points to deplete CENP-B. Data are shown in panel B. **(B)** Representative images of GFP-ZNF518B-transfected cells quantified in panels C and D. Localization of GFP-ZNF518B (green) relative to centromeres in CENP-B^ON^, CENP-B^OFF^, and CENP-B^4h-OFF^ cells. CENP-A (red), CENP-B (blue), and DAPI staining are shown. Areas marked by dashed boxes in the left images are magnified in the right panel. **(C)** Quantification of the fluorescence intensities of nuclear CENP-B signals in CENP-B^ON^, CENP-B^OFF^, and CENP-B^4h-OFF^ cells based on the images in panel B. \*\*\**p* < 0.005, red bars indicate medians. **(D)** Quantification of the fluorescence intensities of GFP-ZNF518B signals at the pericentromeres in CENP-B^ON^, CENP-B^OFF^, and CENP-B^4h-OFF^ cells based on the images shown in panel B. The y-axis shows the relative fluorescence intensity of GFP-ZNF518B signals associated with pericentromeres to total GFP signals in the nucleus. Circles represent measurements from the individual cells shown in panel B, red bars indicate medians. \*\**p* < 0.05. **(E)** Percentage of GFP-ZNF518B foci co-localized with centromeres (CENP-A foci) in CENP-B^ON^, CENP-B^OFF^, and CENP-B^4h-OFF^ cells based on the images in panel B. **(F)** Schematic diagram of the FMIT assay for investigating the interaction between ZNF518B and CENP-B. The CENP-B^133–599^ derivative was used as it cannot bind to the α-satellite repeats at pericentromeres by itself. **(G)** FMIT assay to quantify the co-localization between TetR-EYFP-ZNF518 and Halo-CENP-B^133–599^. Numbers and colors reflect % frequencies of Halo-CENP-B^133–599^ recruitment onto TetR-EYFP-ZNF518B foci. Red shaded boxes represent the ZNF518B segment predicted to be involved in the interaction with CENP-B^133–599^. **(H)** Model depicting the central roles of ZNF518A and ZNF518B in pericentromeric heterochromatin formation via their interactions with CENP-B, HP1, and G9a. The red ellipse and arrow highlight the reaction step focused on in the present figure.

### Interaction between ZNF518B and CENP-B

We further examined the interaction between ZNF518B and CENP-B using the FMIT assay. For this, we constructed Halo-CENP-B^133–599^, which is devoid of the helix-turn-helix domain and thus cannot bind to α-satellites at centromeres.^17^ Halo-CENP-B^133-599^ and TetR-EYFP-ZNF518B (WT and derivatives) were co-expressed in HeLa-03-Int cells (Fig. 4F). As expected, Halo-CENP-B^133–599^ did not localize to the centromeres. Therefore, CENP-B^133–599^ foci reflected the CENP-B recruitment to alphoid*^tetO^* by TetR-EYFP-ZNF518B.

CENP-B^133–599^ was recruited to alphoid*^tetO^* in 26% of cells expressing ZNF518B^WT^, but not in cells expressing TetR-EYFP and lacking ZNF518B. Moreover, ZNF518B^WT^ was recruited to alphoid*^tetO^* in cells expressing TetR:EYFP:CENP-B^133–599^ (Fig. S6B). These results were consistent with the interaction between ZNF518B and CENP-B (Fig. 4G). CENP-B^133–599^ was barely recruited to alphoid*^tetO^* by ZNF518B^1–398^, ZNF518B^1–628^, and ZNF518B^344–628^, suggesting that the ZNF518B segment from aa 629 to the C-terminus is important for the interaction. CENP-B^133–599^ was recruited to alphoid*^tetO^*at high frequencies in cells expressing ZNF518B^344–1074^, ZNF518B^344–987^, ZNF518B^344–877^, ZNF518B^555–1074^, ZNF518B^555–877^, ZNF518B^770–987^, and ZNF518B^770–877^, all of which contain residues 770–877 (Fig. 4G). The recruitment frequency of CENP-B^133–599^ was reduced by 20% in cells expressing ZNF518^344–769^ versus those expressing ZNF518^344–877^ and by 39% in cells expressing ZNF518^555–769^ versus those expressing ZNF518^555–877^. ZNF518B^878–1074^ and ZNF518B^878–987^ did not exhibit CENP-B^133–599^ recruitment. These results support the importance of residues 770–877 of ZNF518B for interaction with CENP-B (Fig. 4H).

We next examined the recruitment of CENP-B to alphoid*^tetO^* in cells expressing ZNF518B^KSS/N^, ZNF518B^KSS/C^, ZNF518B^KSS/NC^, and ZNF518B^770–987^ ^KSS/C^ and carrying substitutions of the PxV(I)xL motifs with KxSxS (Fig. 4G). The PxV(I)xL substitutions generally increased CENP-B recruitment to the *tetO* repeats, implying an inhibitory role of the PxV(I)xL motifs in the interaction between ZNF518B and CENP-B. The derivatives with the N-terminal region deleted showed considerably stronger interactions with both HP1 and CENP-B than full-length ZNF518B (Figs. 3C and 4G). Therefore, we speculate that the interaction domain in some ZNF518 segments is exposed on the protein surface, allowing for easier interactions with the binding partner.

### The C-terminus of ZNF518B interacts with G9a

Proteomics data have revealed an interaction between ZNF518B and G9a (EHMT2).^32^ We attempted to identify the ZNF518B segments that interact with G9a and vice versa using the FMIT assay with cells co-expressing EYFP-TetR-ZNF518 and Halo-G9a. G9a was recruited to *tetO* repeats in control cells expressing ZNF518B^WT^ (Fig. S6D and 5A). G9a was recruited to alphoid*^tetO^* at high frequencies in cells expressing ZNF518B^KSS/C^, ZNF518B^KSS/NC^, ZNF518B^344–1074^, ZNF518B^555–1074^, and ZNF518B^878–1074^, all of which carried the C-terminus, but barely in cells expressing the other ZNF518B derivatives (Fig. 5A). Therefore, the C-terminus (residues 988–1074) of ZNF518B is crucial for interaction with G9a.

**Figure 5.**
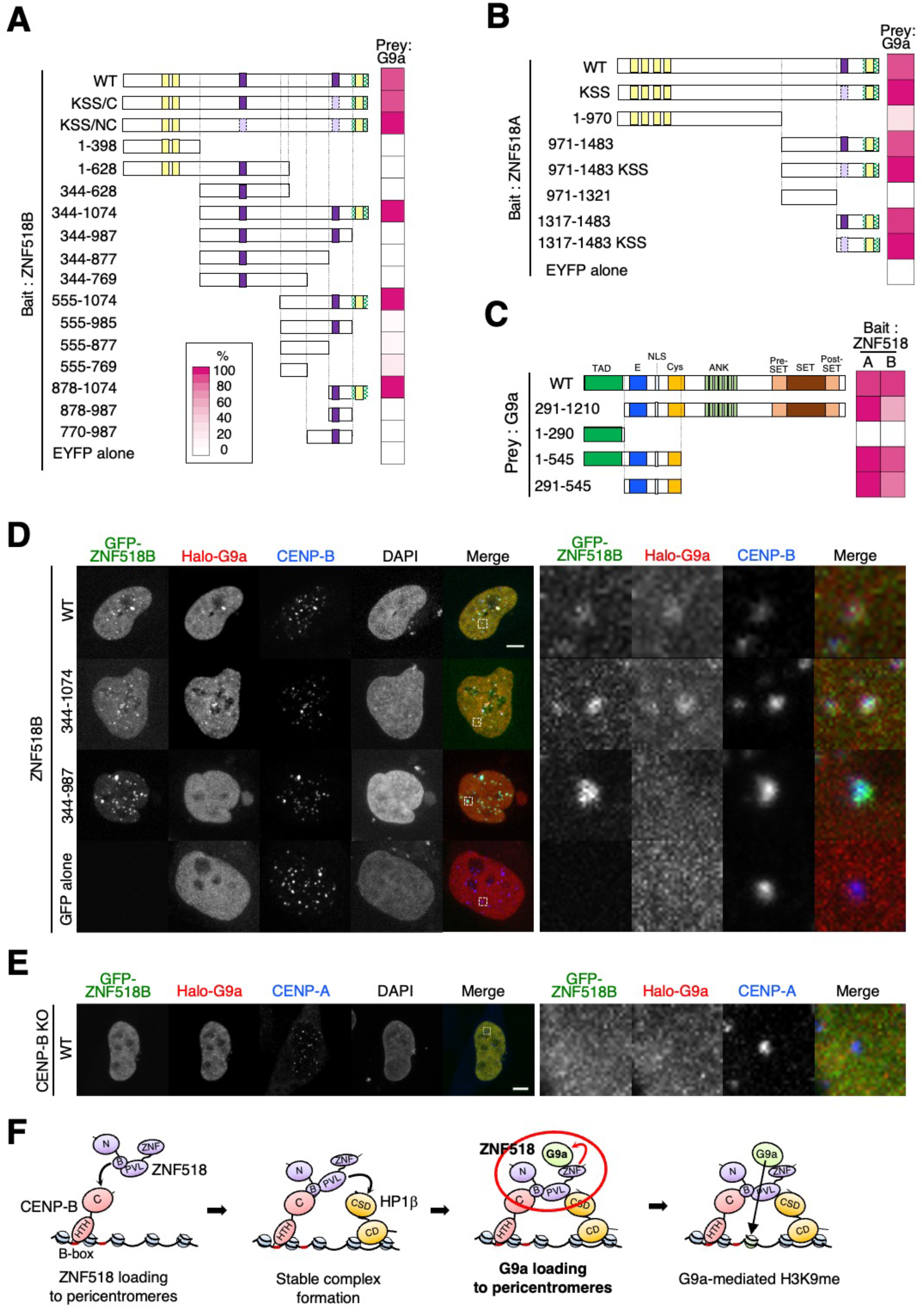
Dissection of ZNF518A and ZNF518B in terms of their requirement for the loading of G9a to centromeres. **(A)** FMIT assay to quantify the co-localization between TetR-EYFP-ZNF518 (WT and 16 derivatives) and Halo-G9a. Green shaded boxes represent the ZNF518B segment potentially involved in its interaction with G9a. **(B)** FMIT assay to quantify the co-localization between TetR-EYFP-ZNF518A (WT and seven derivatives) and Halo-G9a. Green shaded boxes represent the ZNF518A segment potentially involved in its interaction with G9a. **(C)** FMIT assay to quantify the co-localization between Halo-G9a (WT and four derivatives) and TetR-EYFP-ZNF518A or TetR-EYFP-ZNF518B. **(D)** Co-visualization of Halo-G9a (red) and GFP-ZNF518B (WT and two derivatives) or GFP alone (green) in WT cells. Areas marked by dashed boxes in the left images are magnified in the right panel. Scale bar, 5 μm. **(E)** Localization of Halo-G9a (red) and GFP-ZNF518B^WT^ (green) in *CENPB*-KO cells. Scale bar, 5 μm. **(F)** Model depicting the central roles of ZNF518A and ZNF518B in pericentromeric heterochromatin formation via their interactions with CENP-B, HP1, and G9a (see Discussion for details).

We also examined the interaction between ZNF518A and G9a using the FMIT assay. G9a was recruited to alphoid*^tetO^* in cells expressing ZNF518A^WT^, ZNF518A^KSS^, ZNF518A^971–1483^, ZNF518A^971–1483^ KSS, ZNF518A^1317–1487^, and ZNF518A^1317–1483^ KSS, all of which included the C-terminus, but less efficiently in cells expressing ZNF518A^1–970^ and ZNF518A^971–1321^ (Fig. 5B). These results suggested that the C-terminus (residues 1317–1483) of ZNF518A is important for interaction with G9a.

### The segment downstream of the TAD in G9a interacts with ZNF518A/B

G9a and the C2H2-type zinc-finger motifs in ZNF644 and ZNF803, known as WIZ-containing proteins, can interact.^31, 44^ The zinc-finger motifs in WIZ-containing proteins also interact with the transcriptional regulatory CCAAT/enhancer-binding protein-β transactivation domain (TAD) in the N-termini of G9a and GLP.^45^ Because zinc-finger motifs that interact with G9a are highly conserved in the ZNF518A/B C-termini in all species, except *Xenopus laevis* (Fig. S6E), we hypothesized that the C-termini of ZNF518A/B may interact with the G9a TAD.

To test this hypothesis, we conducted FMIT assays with several G9a derivatives (Fig. 5C). In contrast to our hypothesis, G9a^1–290^ was not recruited to alphoid*^tetO^* by ZNF518A/B, whereas G9a^291–1210^, lacking the TAD, was. Moreover, two 291–545-aa segment-containing G9a derivatives, G9a^1–545^ and G9a^291–545^, were recruited to alphoid*^tetO^* by ZNF518A/B. These results indicated that residues 291–545 of G9a, not the TAD, are sufficient for interactions with ZNF518A/B.

### ZNF518B recruits G9a H3K9 methyltransferase to centromeres

Finally, we assessed whether ZNF518 recruits G9a to centromeres. Plasmids expressing GFP-ZNF518B or ZNF518B derivatives and Halo-G9a were co-transfected into U2OS cells, and the localization of Halo-G9a was investigated using the FMIT assay (Fig. 5D). Halo-G9a localized to the centromeres only when expressed with WT or ZNF518B^344–1074^ carrying the C-terminus, not when expressed with ZNF518B^344–987^ lacking the C-terminal zinc-finger (Fig. 5D). Halo-G9a did not localize to the centromeres when GFP-ZNF518B and Halo-G9a were expressed in *CENPB*-KO cells (Fig. 5E). Therefore, CENP-B recruits ZNF518B to centromeres, and ZNF518B in turn recruits G9a H3K9 methyltransferase (Fig. 5F).

## Discussion

This study showed that the 770–877-aa segment in ZNF518B interacts with CENP-B and that CENP-B recruits ZNF518B to centromeres and pericentromeres. Furthermore, the 555–765 segment and PxIxL motif in ZNF518B are required for interaction with the chromo-shadow domain of HP1β. The pericentromere forms constitutive heterochromatin, promoting HP1 accumulation.^8, 9,11^ Therefore, among the ZNF518 proteins recruited to the centromere/pericentromere by CENP-B, those assembled in the pericentromere are likely to maintain a stable association with the chromatin through interaction with HP1. The ZNF518B C-terminus carrying the zinc-finger motif interacts with the segment downstream of the TAD in G9a, and CENP-B is required for the centromeric localization of G9a. Based on these findings, we propose the following mechanisms: 1) CENP-B binds to α-satellite DNA to initiate heterochromatin formation; 2) CENP-B recruits ZNF518B to centromeres and pericentromeres; 3) HP1β stabilizes ZNF518 only at pericentromeres; and 4) ZNF518B interacts with G9a for H3K9 methylation at the pericentromeres.

These conclusions do not disagree with a previously proposed mechanism involving SUV39H1 histone H3K9 methyltransferase.^26, 27^ HP1-dependent SUV39H1 recruitment and short RNA-mediated heterochromatinization are important for pericentromeric heterochromatin maintenance.^26, 28^ We suggest that G9a recruited by CENP-B and ZNF518B is required for pericentromeric heterochromatin establishment and that the CENP-B/ZNF518B/HP1β/G9a axis targets heterochromatin factors to pericentromeres, where specificity is achieved by the binding of CENP-B to CENP-B box sequences in pericentromeric alphoid repeats.

ZNF518A is also recruited to pericentromeres in a CENP-B-dependent manner (Fig. S1) and interacts with HP1 and G9a (Figs. 5, S5). ZNF518A and ZNF518B are evolutionarily conserved in all vertebrates higher than amphibians. Therefore, we speculate that pericentromeric heterochromatin formation by ZNF518 is conserved among many vertebrates. Although further investigation is needed, our study provided insights into the potential mechanisms underlying the targeting of heterochromatin to human pericentromeres and indicated that a similar mechanism may exist in many species.

## Material and Methods

### DNA constructs

DNA fragments encoding human ZNF518A, ZNF518B, CENP-B^133–599^, HP1α, HP1β, HP1γ, and G9a were PCR-amplified (primers are listed in Table S1) from a cDNA library generated by the Kazusa ORFeome Project and Genome Network Project Clones by RIKEN BRC. The fragments were inserted into an entry vector (pENTR™ 4 Dual Selection Vector; #A10465, Thermo Fisher Scientific) between the *Xmn*I (#R0194S, New England BioLabs [NEB]) and *EcoR*V (#R0195S, NEB) sites using the In-Fusion HD Cloning Kit (#Z9648N, Takara Bio), yielding entry plasmids pENTR4XE153, pENTR4XE127, pENTR4τιCENPB, pENTR4HP1α, pENTR4HP1β, pENTR4HP1γ, and pENTR4G9a, which comprised the open reading frames encoding the above respective proteins.^46–50^

DNA fragments encoding ZNF518A derivatives (ZNF518A^KSS^, ZNF518A^1–970^, ZNF518A^971–1483^, ZNF518A^971–1483KSS,^ ZNF518A^971-1321^, ZNF518A^1317–1483^, and ZNF518A^1317–1483KSS^), ZNF518B derivatives (ZNF518B^KSS/N^, ZNF518B^KSS/C^, ZNF518B^KSS/NC^, ZNF518B^1–398^, ZNF518B^1-628^, ZNF518B^344–628^, ZNF518B^344–1074^, ZNF518B^344–987^, ZNF518B^344-877^, ZNF518B^344-769^, ZNF518B^555–985^, ZNF518B^555–877^, ZNF518B^555–769^, ZNF518B^878v1074^, ZNF518B^878–987^, ZNF518B^770–987^, ZNF518B^770–987KSS/C^, and ZNF518B^770–877^), HP1β derivatives (HP1β^N(1–92)^ and HP1β^C(93–185)^), and G9a derivatives (G9a^1–92^ and G9a^93–185^) were PCR-amplified from pENTR4XN153, pENTR4XN127, pENTR4HP1β, and pENTR4G9a, respectively, using the primers listed in Table S1.

To overexpress the target proteins in human or *E. coli* cells, plasmids were generated by DNA recombination between the above-mentioned entry plasmids and pDEST131NGFP (for EGFP fusion proteins),^51^ pDEST152JTY (for EYFP-TetR fusion proteins), pDEST155Halo (for Halo-tagged proteins), pDEST174HATMBP (for HAT [KDHLIHNVHKEEHAHNK]-MBP fusion proteins),^52^ or pDEST172HGST (for His_6_-GST fusion proteins) using Gateway LR Clonase II Enzyme mix (#11791020, Thermo Fisher Scientific).

For CRISPR-mediated double-strand break site formation, 54-bp DNA fragments were generated by annealing two complementary single-stranded oligonucleotides. The fragments were inserted into the *Age*I (#R3552S, NEB) site of pTORA14 using the In-Fusion HD Cloning Kit,^53^ yielding pTORA14AAVS1, pTORA14CENP-B_C, pTORA14HA4, pTORA14HB7, and pTORA14HG1, which were used to induce DNA cleavage at adeno-associated virus integration site 1 in *CENPB, HP1ALPHA, HP1BETA*, and *HP1GAMMA*, respectively. The DNA sequences targeted by the guide RNAs are listed in Table S2.

To construct the plasmid directing homologous recombination (HR) at the 3′ end of *CENPB*, a DNA fragment was PCR-amplified from human genomic DNA using Tks Gflex DNA Polymerase (#R060A, Takara Bio) and the primers listed in Table S1. This yielded a 584-bp DNA fragment containing the 267-bp left homological arm (HRL) and 314-bp right homological arm (HRR) regions identical to the target locus. The amplified DNA fragment comprising the HRL and HRR regions was cloned into pCR-Blant (#K275020, Thermo Fisher Scientific). To construct HR donor plasmids (pKANA1CENPB and pKANA2CENPB), mAID cassettes containing the mAID-encoding region and a hygromycin or blasticidin resistance gene were PCR-amplified from pMK287 or pMK288 and inserted between the HRL and HRR regions using the In-Fusion HD Cloning Kit.^43^ The resultant plasmids were used to generate a human cell line expressing the CENP-B:mAID fusion protein.

### Cell lines

HeLa-Int-03 cells containing approximately 15,000 repeats of the *tetO* sequence in chromosome 21 with and without *CENPB* or *HP1* KO were established as previously described.^17,37,54^ A U2OS-*os*TIR1 cell line, which is the parental cell line of *CENPB*-cKO cells, was generated by transfecting U2OS cells with pTORA14AAVS1 and pMK243 using Lipofectamine LTX Reagent (#A12621, Thermo Fisher Scientific)^43^ followed by puromycin selection (1 µg/mL, #10-2100, Focus Biomolecules). *CENPB-*cKO cells were generated by transfecting the U2OS-*os*TIR1 cells with pTORA14CENP-B_C, pKANA1CENPB, and pKANA2CENPB followed by selection on hygromycin B (1 mg/mL, #089-06151, Wako) and blasticidin S (5 µg/mL, #ant-bl-05, InvivoGen).

### Cell culture

U2OS and HeLa cells were cultured in Dulbecco’s modified Eagle’s medium (DMEM; #044-29765, FUJIFILM Wako) supplemented with 10% (v/v) fetal bovine serum (Bovogen Biologicals), 100 U/mL of penicillin, and 100 µg/mL of streptomycin (#168-23191, Wako) at 37 °C in a humidified incubator with 5% CO_2_. To generate CENP-B^OFF^ cells, *CENPB*-cKO cells were cultured with 0.2 µg/mL Dox (#D3447, Sigma-Aldrich) for 24 h and subsequently with 0.5 mM IAA (#094-07121, Wako) for 24 h before use.

### Indirect immunofluorescence and microscopy

U2OS and HeLa cells in the exponential growth phase were seeded onto coverslips at 30–40% confluence and allowed to grow overnight. To overexpress EGFP-ZNF518A and EGFP-ZNF518B for indirect immunofluorescence experiments, pDEST131NEGFP153 and pDEST131NEGFP127, respectively, were transfected into the cells using Lipofectamine LTX. Twenty-four hours later, the cells were fixed with 4% (v/v) paraformaldehyde (#163-20145, Wako) in phosphate-buffered saline (PBS) for 10 min and permeabilized with 0.15% (v/v) Triton X-100 (#1610407, Bio-Rad) in PBS for 2 min. The cells were treated with 1% (v/v) bovine serum albumin (BSA; #A7284, Sigma-Aldrich) in PBS and subsequently incubated with primary antibodies, including mouse anti-CENP-A (1:200 dilution, #ab13939, Abcam), mouse anti-CENP-B (1:50,000, 5E6C1),^16^ rabbit anti-CENP-B (1:200, #ab25734, Abcam), 1:600-diluted rabbit anti-CENP-C (#554),^55^ mouse anti-HP1α (1:1,000, #MAB3584, Merck Millipore), rabbit anti-HP1β (1:800, #8676 D2F2, Cell Signaling Technology), mouse anti-HP1γ (1:3,000, #MAB3450 2MOD-1G6, Merck Millipore), or rabbit anti-histone H3K9me3 (1:1,000, #ab8898, Abcam). The cells were washed with PBS thrice for 5 min and incubated with Alexa-conjugated secondary antibodies (1:600). DNA was stained with 0.1 µg/mL DAPI (#H-1200, Vector Laboratories).

The cells were observed using an Olympus FV1000 confocal microscope with a UPlanSApo 60×/1.35 oil-immersion objective lens (Olympus) and FV10-SAW2.1 software (Olympus). Images were also acquired using a Nikon Confocal A1Rsi microscope with a CFI Plan Apochromat VC 60×/1.40 oil-immersion objective lens (Nikon) and FNIS-Elements AR software (Nikon). The images were acquired at 0.5-μm intervals in the z-axis and Kalman-filtered to suppress background noise.

### FMIT

pDEST172JTY and pDEST155Halo encoding EYFP-TetR and Halo fusion proteins, respectively, were transfected into HeLa cells carrying the *tetO* array at 60% confluence using Lipofectamine 3000 (#L3000001, Thermo Fisher Scientific). Twenty-four hours later, the cells were incubated with 1 nM Halo Tag TMR Ligand (G825A, Promega) at 37 °C for 2 h. The labeled cells were washed with DMEM and incubated in DMEM at 37 °C for 30 min. Then, the cells were fixed with 4% (v/v) paraformaldehyde in PBS for 10 min and DNA was stained with 0.1 µg/mL DAPI. EYFP and TMR signals were observed using an Olympus BX53 fluorescence microscope with a UPlanSApo 60×/1.35 oil-immersion objective lens and cellSens software (Olympus). More than 100 cells were analyzed for each sample.

### Quantification of microscopic data

Fluorescence quantitation and line plot analyses of the fluorescence signals were performed using ImageJ 1.50e.^56^ To quantitate fluorescence, z-stack images were captured at every 500 nm between the lower and upper edges of nuclei and merged using the maximum intensity z-projection function in ImageJ. In projected images, nuclei and centromeres were defined by DAPI and CENP-A signals, respectively. Nucleus-and centromere-associated GFP signals in the cells were estimated, and their ratio was calculated to determine whether GFP-fused proteins were enriched at centromeres. Statistical analyses were performed using R v.3.5.1 (R Foundation for Statistical Computing, http://www.r-project.org).

For the line plot analyses, the slice showing the strongest signal of a centromeric protein (CENP-A or CENP-B) and GFP-ZNF518 was selected from the z-stack images. Using the RGB plot profile function in ImageJ, signal intensities of the centromeric protein and GFP-ZNF518 were quantified for each pixel along the indicated lines.

### Protein purification

His_6_-GST-HP1 and HAT-MBP-ZNF518B^770–987^ were purified from BL21 *E. coli* cells. To express the target proteins, *E. coli* cells carrying pDEST173HGSTHP1 for His_6_-GST-HP1 or pDEST174HATMBP127 for HAT-MBP-ZNF518B^770-987^ were cultured in L broth supplemented with 0.1 mM IPTG (#096-05143, Wako) at 30 °C for 2 h. His_6_-GST-HP1 proteins were extracted using a buffer containing 50 mM Tris-HCl (pH 8.0; #201-06273, Wako), 500 mM NaCl (#191-01665, Wako), 10 mM β-mercaptoethanol (#131-14571, Wako), and 10% glycerol (#075-00611, Wako). HAT-MBP-ZNF518B^770-987^ was extracted using above buffer including 1% Tween 20 (#H5152, Promega). The target proteins were affinity-purified using Ni-NTA agarose (#143-09763, Wako).

### MBP PD

All MBP PD procedures were performed using PD buffer comprising 25 mM Tris-HCl (pH 8.0), 100 mM NaCl, 1 mM EDTA (#E5134, Sigma-Aldrich), 1 µM DTT (#GE17-1318-02, Sigma-Aldrich), 1% Tween 20, 10% glycerol, and 0.02% BSA. First, 2 µg HAT-MBP-ZNF518B^770–987^ protein and 5 µg His_6_-GST-HP1 protein, purified as described above, were mixed and incubated under rotation at 4 °C for 4 h. Next, 50 µg amylose magnetic beads (#E8035S, NEB) was added to the protein mixture, which was further incubated under rotation at 4 °C for 1 h. After washing the beads six times with 1 mL of PD buffer, the sample was boiled with 1× SDS loading buffer at 95 °C for 5 min. The eluate was subjected to immunoblotting.

### Immunoblotting

Immunoblotting was carried out using mouse anti-MBP (1:200, #sc-271524, Santa Cruz Biotechnology), mouse anti-GST (1:200, #sc-138, Santa Cruz Biotechnology), rabbit anti-HP1α (1:1,000, #2616, CST), mouse anti-HP1α (#MAB3584, Merck Millipore), rabbit anti-HP1β (1:1,000,#8676, CST), rabbit anti-HP1γ (1:1,000,#2619, CST), mouse anti-HP1γ (1:1,000, #MAB3450, Merck Millipore), and mouse anti-GAPDH (1:10,000, #5A12,016-25523, Wako) primary antibodies. The target proteins were detected using IRDye 800CW donkey anti-rabbit IgG and anti-mouse IgG (1:15,000, #926-32213, #926-32210, Li-COR Biosciences) secondary antibodies.

### Amino acid sequence analysis

The Basic Local Alignment Search Tool was used to identify amino acid similarities between query sequences and sequences deposited in DDBJ/GenBank/EMBL.^57^ A neighbor-joining phylogenetic tree was constructed using MEGA X software.^58^

## Acknowledgments

The authors thank Iyo Toramoto, Kana Yamazoe, and Fusako Kawasaki (Kochi University) for technical assistance; Koichiro Otake (Kazusa DNA Research Institute) for providing preliminary data; Koichi Honke, Hideaki Kuge, Kaoru Miyahara, Arisa Miyagawa-Yamaguchi, and Tatsuyuki Yamashita (Kochi University) for scientific advice; and William Earnshaw (University of Edinburgh) for critical manuscript reading. This work was supported by grants from JSPS KAKENHI (grant No. 18K06061), the Kato Memorial Bioscience Foundation, and the Takeda Science Foundation to SO, by grants from JST, CREST (grant No. 18070874), and MEXT KAKENHI (grant No. 20H00473), and by a grant from the Kazusa DNA Research Institute Foundation to HM.

## Author contributions

SO, JO, and HM designed the study. SO, JO, and NS performed the experiments. SO, JO, and HM analyzed the data. SO, JO, KN, and HM wrote the manuscript.

