## supplemental information for "Zinc-finger protein 518 plays a crucial role in pericentromeric heterochromatin formation by linking satellite DNA to heterochromatin"

**Supplemental information figure legends and tables**

**Figure S1. CENP-B-dependent localization of ZNF518B at pericentromeres.**

**(A)** Localization of GFP-ZNF518B (green) in mitotic cells. Scale bar, 5 μm.

**(B)** Protein domains present in ZNF518A and ZNF518B. Yellow and purple boxes indicate zinc-finger (ZF) and PxV(I)xL motifs, respectively.

**(C)** Localization of GFP-ZNF518A (green), CENP-A (red), and CENP-B (blue) in WT (upper) and *CENPB*-knockout (KO) cells (lower) with and without overexpression (OX) of CENP-B-$FL_3$.

**(D)** Model depicting the central roles of ZNF518A and ZNF518B in pericentromeric heterochromatin formation via their interactions with CENP-B, HP1, and G9a.

**Figure S2. Dissection of ZNF518A and ZNF518B to identify their segments required for pericentromeric localization.**

**(A)** Quantification of GFP-ZNF518B (WT and 12 derivatives) signals at pericentromeres. Relative fluorescence intensities of GFP-ZNF518B signals associated with pericentromeres to total GFP signals in the nucleus are shown (right). Circles represent measurements from the individual cells shown in Figure 2, red bars indicate medians. ***$p$ < 0.005 versus WT.

**(B)** Localization of the ZNF518A derivatives at pericentromeres. The schematic representation of the respective ZNF518A derivatives is drawn approximately to scale. Yellow boxes indicate zinc-finger motifs. The PxVxL motif (purple boxes) was substituted with KxSxS (light purple boxes), shown as KSS. GFP-fused ZNF518A signals (WT and six derivatives; green) were co-visualized with the centromeric marker

CENP-A or CENP-B (right panels). Images labeled with “A” show CENP-A foci, all other images show CENP-B signals. Areas marked by dashed boxes in the left images are magnified in the right panel. Numbers and colors (left) reflect % frequencies of the cells with GFP-ZNF518A signals at pericentromeres. Scale bar, 5 μm.

**Figure S3. Multiple sequence alignment of ZNF518A and ZNF518B from various species.**

ZNF518A and ZNF518B aa sequences from *Homo sapiens* (Q6AHZ1 and Q9C0D4), *Bos taurus* (E1BG18 and A0A3Q1M9T7), *Mus musculus* (B2RRF6 and B2RRE4), *Gallus* (E1BXZ2 and F1NRI7), *Nipponia nippon* (A0A091UJX0 and A0A091UXH1), *Amazona aestiva* (A0A0Q3M877 and A0A0Q3PD60), *Chelonia mydas* (M7C1U2 and M7BLJ2), *Xenopus laevis* (A0A1L8FJ56, A0A1L8FEE5, and A0A1L8HKU8), *Danio rerio* (U3JAE4), *Maylandia zebra* (A0A3P9BDT7), *Mastacembelus armatus* (A0A3Q3NFI5), *Oryzias latipes* (H2MVV3), *Salmo trutta* (A0A673Y1T4, A0A673XJD3, and A0A673W0W1), and *Scyliorhinus torazame* (A0A401PBJ2, A0A401NXE1, and A0A401PBJ2) were aligned using MEGA X and represented in a phylogenetic tree.[58] The secondary structure of *H. sapiens* ZNF518B was predicted using Jpred (top). ZNF518A/B aa sequences around the PxVxL motifs are shown. The aa sequence and secondary structure of *H. sapiens* CAF-1 (PDB:1DZ1) are shown at the bottom. Background colors of aa residues reflect their biochemical properties. Asterisks indicate residues that are conserved among all species compared.

**Figure S4. Localization of ZNF518B derivatives at heterochromatin domains in *CENPB*-KO cells.**

**(A)** Localization of GFP-tagged ZNF518B$^{WT}$ and ZNF518B$^{770–987}$ to centromeres (CENP-A; red) in *CENPB*-KO cells.

**(B)** Co-localization of GFP-ZNF518B$^{770–987}$ and histone H3K9me3 (red) in *CENPB*-KO cells.

**(C)** Model depicting the central roles of ZNF518A and ZNF518B in pericentromeric heterochromatin formation via their interactions with CENP-B, HP1, and G9a. The red ellipse and arrow highlight the reaction step focused on in the present figure.

**Figure S5. ZNF518A/B interact with HP1s.**

**(A)** Tethering of YFP-ZNF518B derivatives (green) containing a PxIxL motif to the *tetO* repeats results in the accumulation of Halo-HP1β (top; red).

**(B)** Heatmap summarizing the interactions between ZNF518A and heterochromatic factors. Co-localization frequencies of TetR-EYFP-ZNF518A (WT or derivatives) with heterochromatic factors Halo-HP1α, -HP1β, and -HP1γ were determined. Colors reflect the % frequencies of cells exhibiting co-localization between TetR-EYFP-ZNF518A and the heterochromatic factors at the *tetO* repeats.

**(C)** Dissociation timing of TetR-EYFP-ZNF518A from the *tetO* repeats was microscopically assessed. The percentage of transfected cells carrying TetR-EYFP-ZNF518B foci was plotted against Dox treatment time.

**Figure S6. ZNF518A/B interact with CENP-B and G9.**

**(A)** mAID:CENP-B depletion was assessed by immunofluorescence microscopy. CENP-B (green) was visualized in CENP-B$^{ON}$ (a and a′ panels) and CENP-B$^{OFF}$ cells (panels b and b′). CENP-B$^{OFF}$ cells were prepared by Dox treatment for 24 h and subsequent indole-3-acetic acid (IAA) treatment for 24 h.

**(B)** Tethering of TetR:YFP:CENP-B (green) to the *tetO* repeats results in the accumulation of Halo-ZNF518B (top, red) or ZNF518A (bottom, red) at the locus, whereas the tethering of TetR-YFP (green) does not.

**(C)** Heatmap summarizing the interactions between ZNF518A and heterochromatic factors. Co-localization frequencies of TetR-EYFP-ZNF518A (WT or derivatives) with CENP-B$^{133–599}$ were determined. Colors reflect the % frequencies of the cells exhibiting co-localization between TetR-EYFP-ZNF518A and the heterochromatic factors at the *tetO* repeats.

**(D)** Schematic diagram of the FMIT assay for investigating the interaction between ZNF518B and G9a. *CENPB*-KO cells were used.

**(E)** Multiple sequence alignment of the C2H2-type zinc-finger domains in the C-termini of ZNF518A and ZNF518B from various species. Background colors of aa residues reflect their biochemical properties. Asterisks and colons indicate residues that are conserved among all species compared.

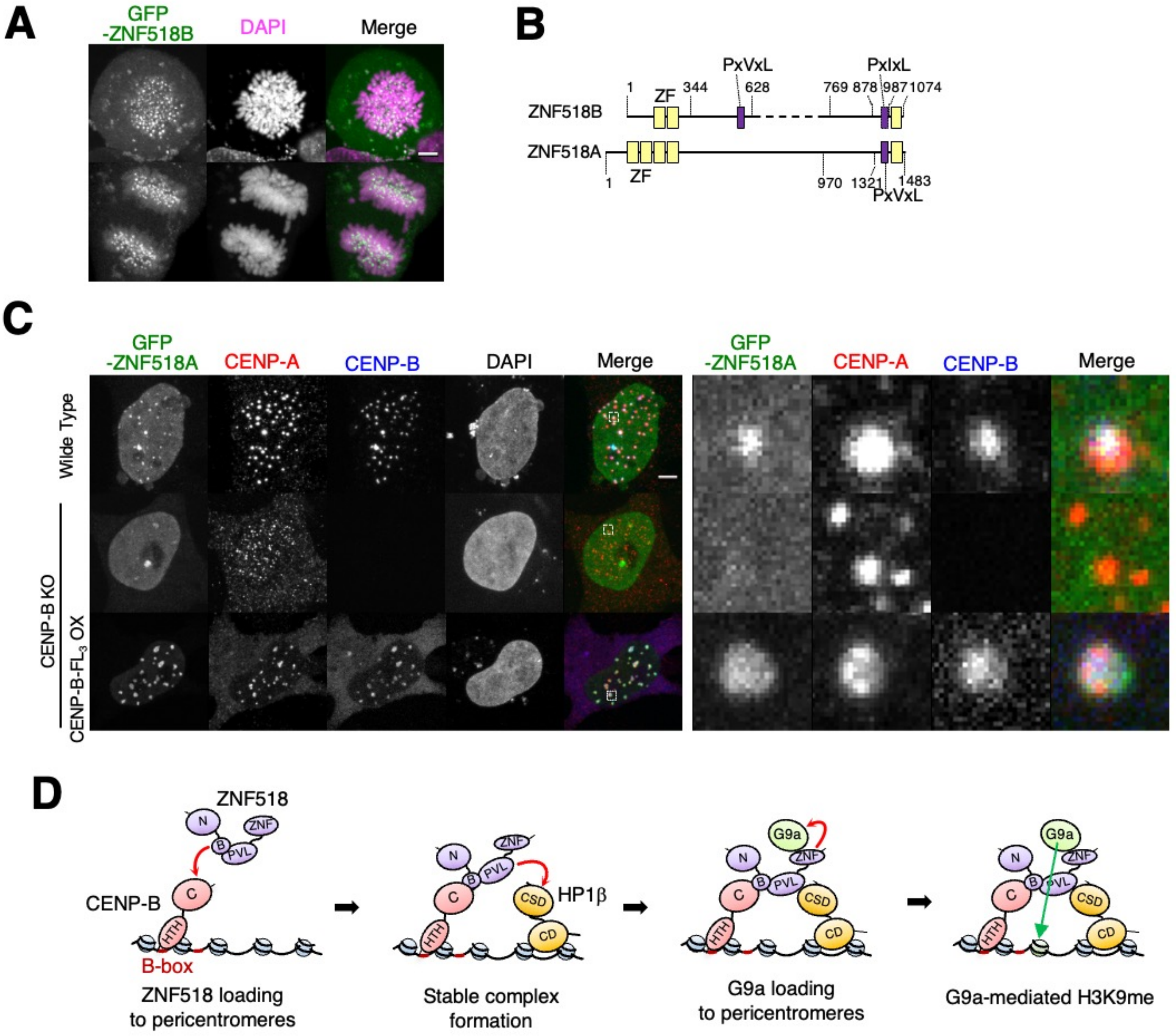
A
GFP
-ZNF518B
DAPI
Merge
B
PxVxL
PxIxL
1
344
628
769 878
987
1074
ZF
ZNF518B
ZNF518A
ZF
1
970
1321
1483
PxVxL
C
GFP
-ZNF518A
CENP-A
CENP-B
DAPI
Merge
GFP
-ZNF518A
CENP-A
CENP-B
Merge
Wide Type
CENP-B KO
CENP-B-FL3 OX
D
ZNF518
N
ZNF
B
PVL
CENP-B
C
HTH
B-box
ZNF518 loading
to pericentromeres
HP1β
CSD
CD
Stable complex
formation
G9a
G9a loading
to pericentromeres
G9a-mediated H3K9me

Ohta *et al.*, Fig. S2

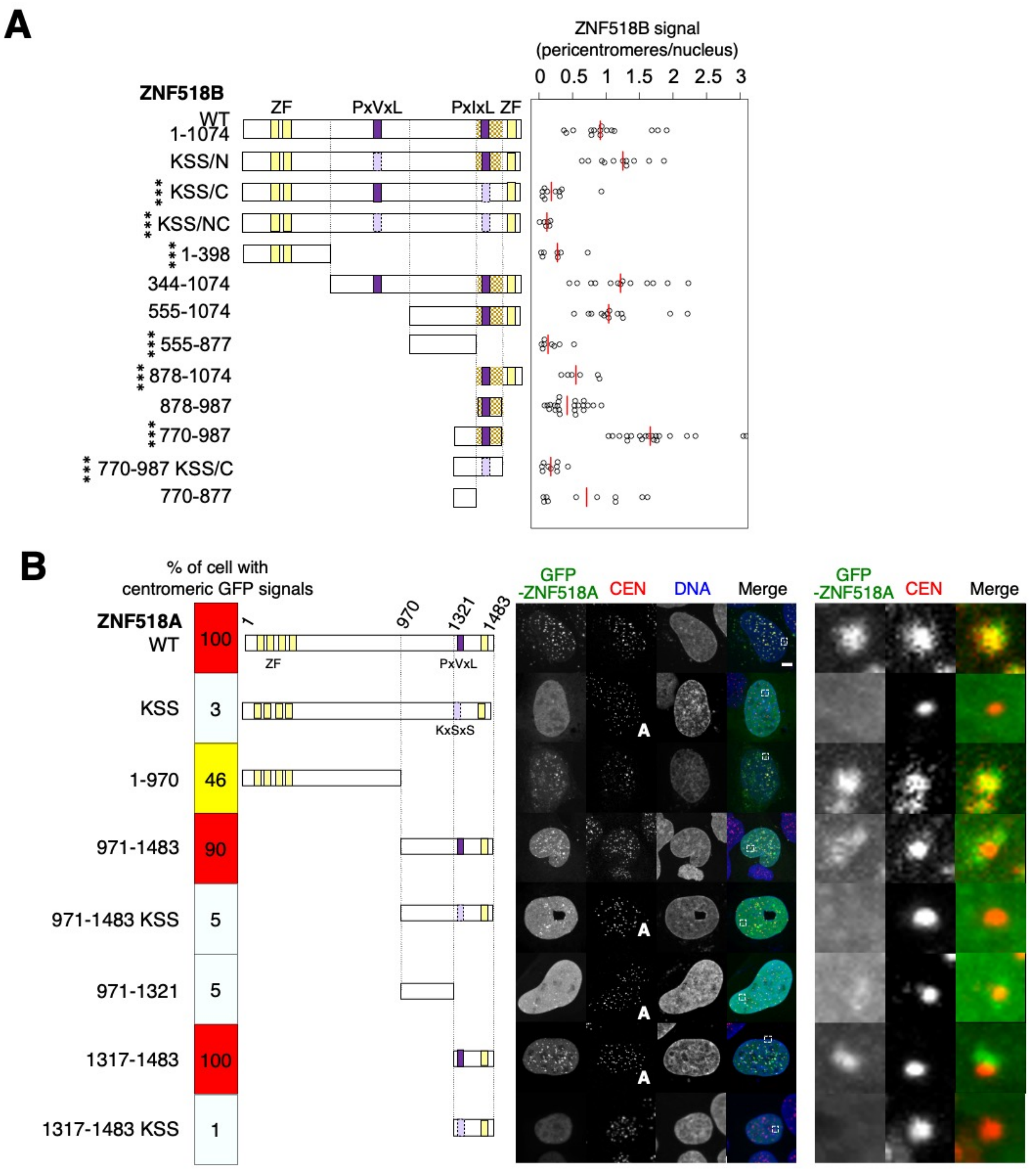

Ohta *et al.*, Fig. S3

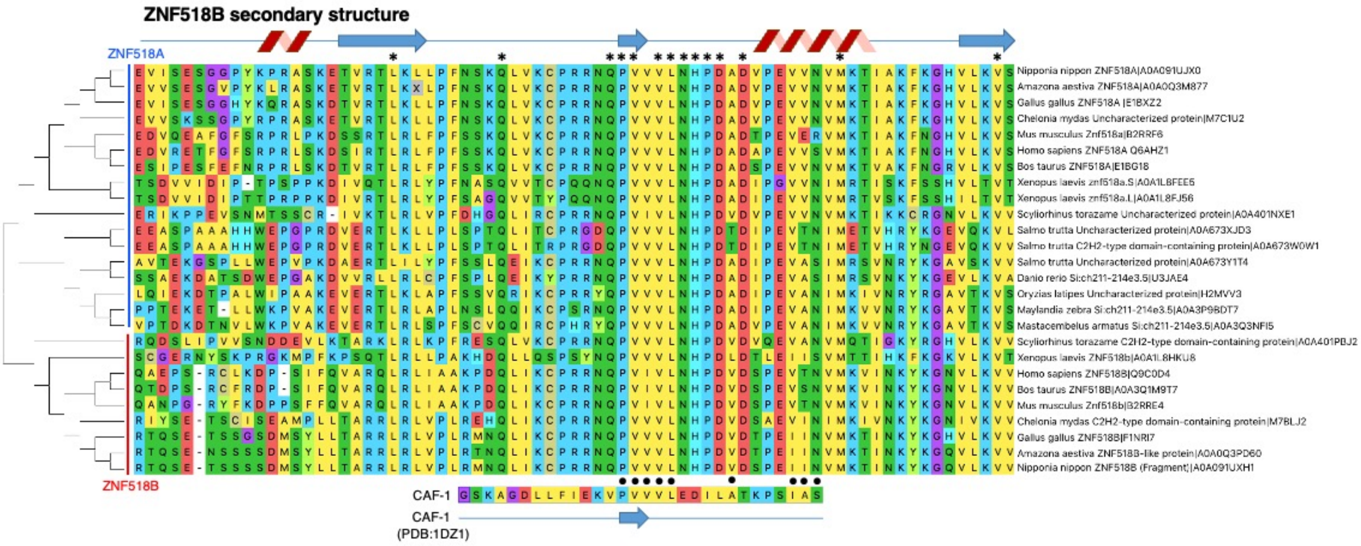

Ohta *et al.*, Fig. S4

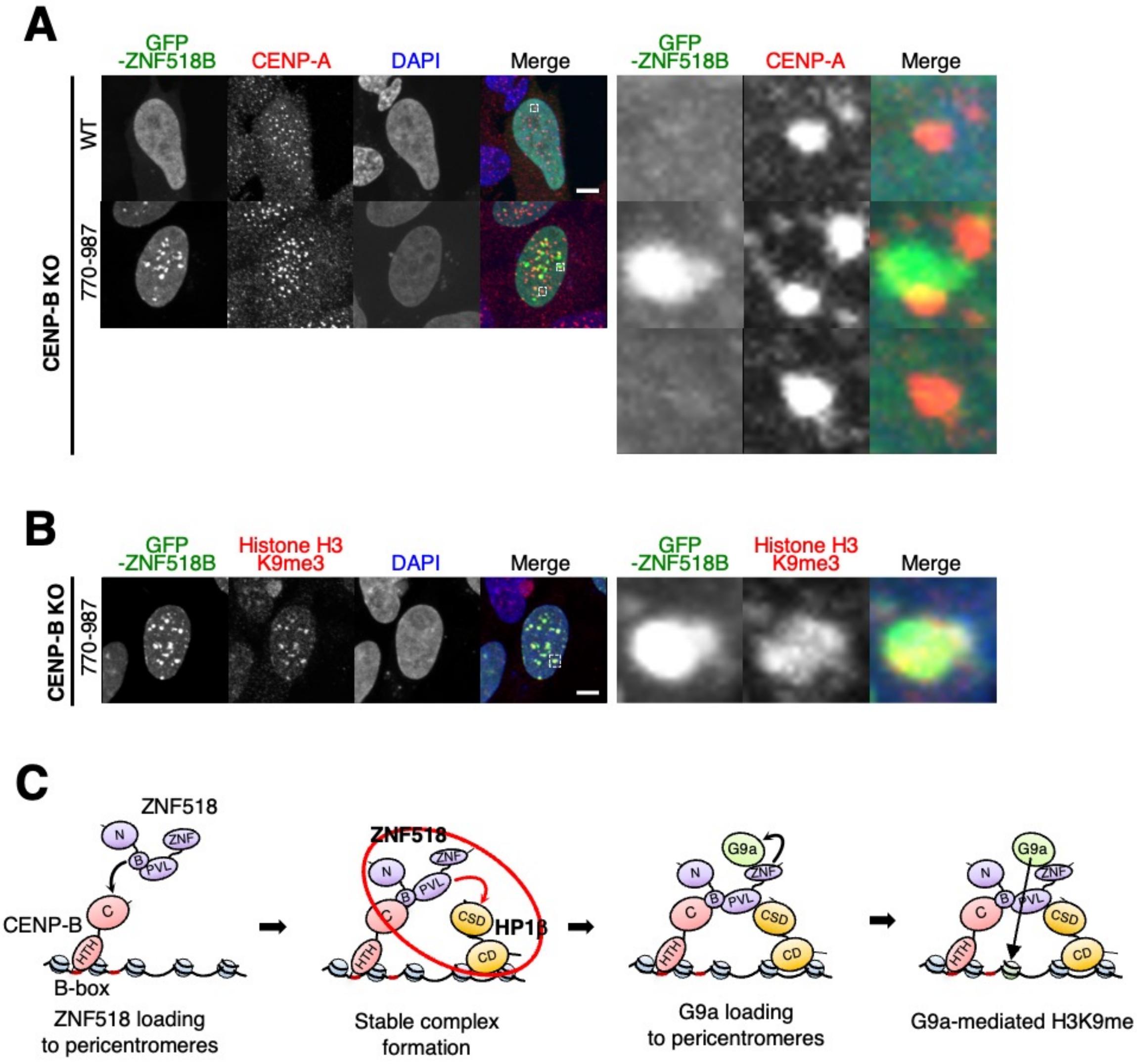

Ohta *et al.*, Fig. S5

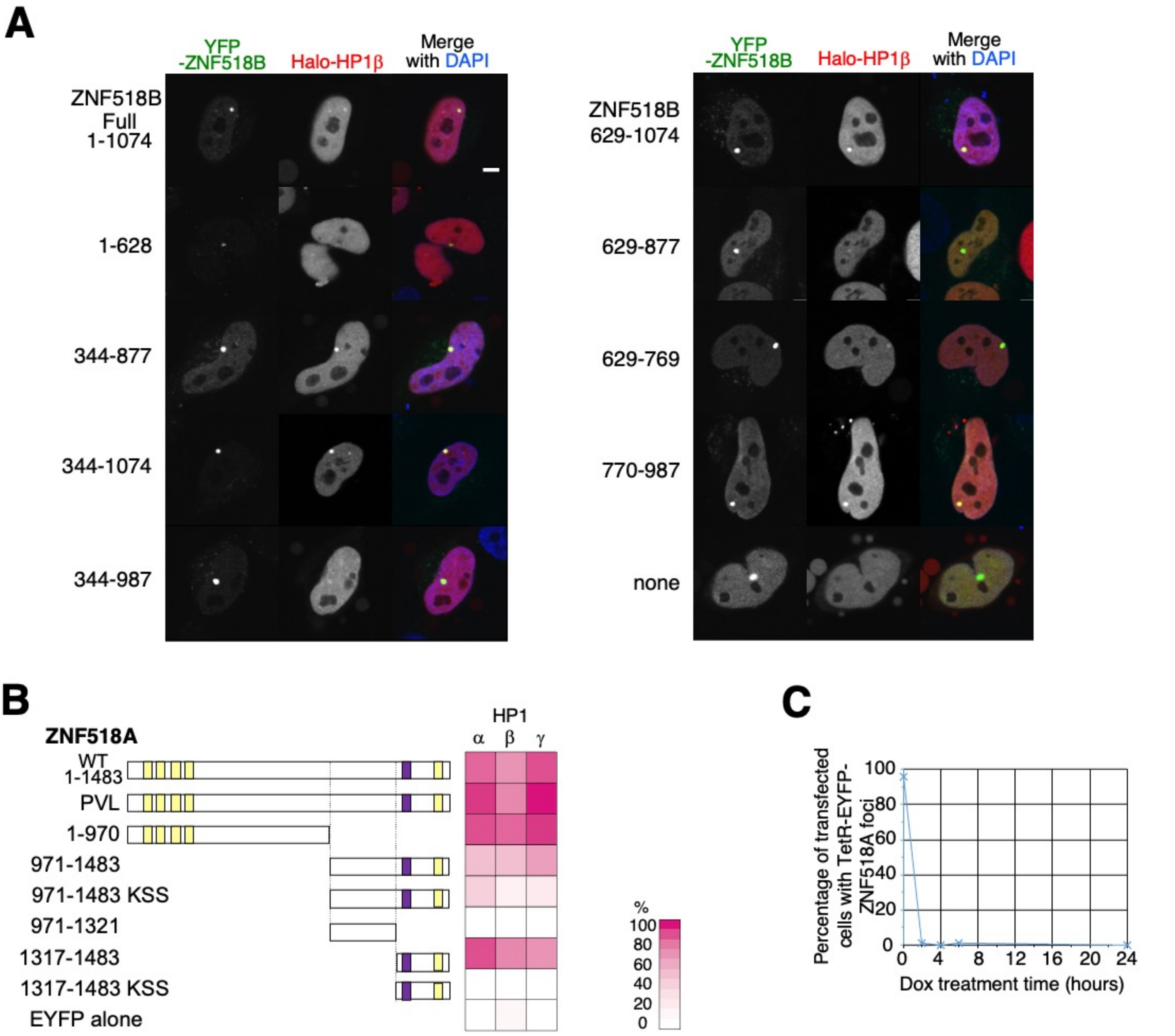

Ohta *et al.*, Fig. S6

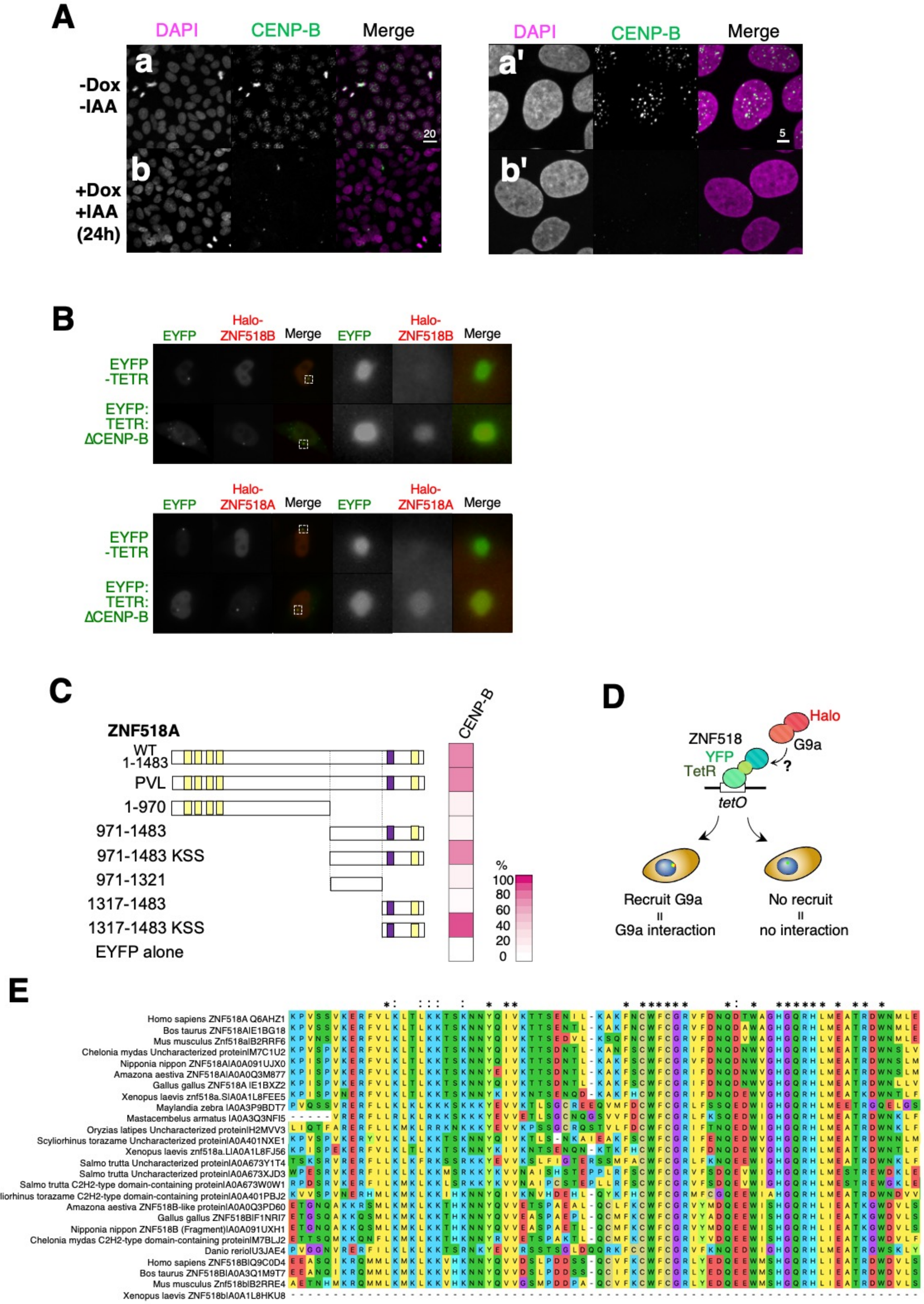

**Supplemental Table S1.** Sequences of the primers used in this study.

| Primer | Sequence (5' → 3') |
|---|---|
| o-Z518A_FW_pE4XmnI | TCCACCATGGGAACCATGCCATCTGAACAGAAACAG |
| o-Z518A_RV_pE4EcoRV | AAGCTGGGTCTAGATGTTCTAACATGTTCCAATCCC |
| o-Z518A_PxVxLmut_FW | AAAAAGTCAGTGTGTCAAATCATCCTGACGCAGATG |
| o-Z518A_PxVxLmut_RV | TGACACACTGACTTTTTGGTTTCTCCTAGGACATTTC |
| o-Z518A_970_upper | GGTATCCCTGCTTCTCTTCATCTAGACCCAGCTTTCTT |
| o-Z518A_970_lower | AAGAGAAGCAGGGATACCTTGTGAATTAACCAATGG |
| o-Z518A_971_upper | GAAACCTTTGTAAACAAGAAACCTGGGATGGTTTTAACAC |
| o-Z518A_971_lower | TTGTTTACAAAGGTTCCCATGGTGGAGCCTGCTTTTTTGT |
| o-Z518A_1317_upper | CCTAGGCTTTCAAAAGATTCCATCAGAACTTTGCG |
| o-Z518A_1317_lower | CTTTTGAAAGCCTAGGGGTTCCCATGGTGGAGCCTGCTTTTTTG |
| o-Z518A_1321_upper | ACCTAGGCTTTCAAAACATCTAGACCCAGCTTTCTT |
| o-Z518A_1321_lower | TTTTGAAAGCCTAGGTCTGCTAAATCC |
| o-Z518B_FW_pE4XmnI | TCCACCATGGGAACCATGACCATTGCTACATGTGC |
| o-Z518B_RV_pE4EcoRV | AAGCTGGGTCTAGATGTTTGCCCTTGGAGGAAAGAAC |
| o-Z518B_PxVxLmut_FW | AAGTCCTCTGCCAAAAAAGGTCATTCTCTAGCATCTG |
| o-Z518B_PxVxLmut_RV | CCTTTTTTGGCAGAGGACTTAAAAGCAGTTCTATGTGG |
| o-Z518B_PxIxLmut_FW | AAAGTCAGTGTGTCAAACCACCCTGATGTGGAC |
| o-Z518B_PxIxLmut_RV | TTGACACACTGACTTTCTGGTTCCGACGGGGAC |
| o-Z518B_398_upper | GTGTGCACAACAATGGAAAACATCTAGACCCAGCTTTCTT |
| o-Z518B_398_lower | TTTTCCATTGTTGTGCACACTAGAATTTGG |
| o-Z518B_628_upper | GTGGGAAGCCGTCATCTTTGATCTAGACCCAGCTTTCTT |

| | | |
|---|---|---|
| 1 | o-Z518B_628_lower | CAAAGATGACGGCTTCCCACAGGACTCAATTAACTGAGC |
| 2 | o-Z518B_344_upper | GCTCCACACTTTCTGCAGAAAAAACAAAGTC |
| 3 | o-Z518B_344_lower | CTGCAGAAAGTGTGGAGCCTGCTTTTTTGT |
| 4 | o-Z518B_987_upper | TATTTAAGTGCTGGTTTTGTCATCTAGACCCAGCTTTCTT |
| 5 | o-Z518B_987_lower | ACAAAACCAGCACTTAAATACACACTGTGAAG |
| 6 | o-Z518B_887_upper | TGATAGCAGCTAAGCCAGATCATCTAGACCCAGCTTTCTT |
| 7 | o-Z518B_877_lower | ATCTGGCTTAGCTGCTATCAGTCTTAG |
| 8 | o-Z518B_769_upper | GACAGGCGGACTCAGACTTACATCTAGACCCAGCTTTCTT |
| 9 | o-Z518B_769_lower | TAAGTCTGAGTCCGCCTGTCTCACAGGGC |
| 10 | o-Z518B_878_upper | CCCAGTTGATTAAGTGTCCCCGTCGG |
| 11 | o-Z518B_878_lower | CACTTAATCAACTGGGAGCCTGCTTTTTTGT |
| 12 | o-Z518B_770_upper | CAGCCTTTAAGAAGTGAAAGGGGGCCAATAG |
| 13 | o-Z518B_770_lower | CTTTCACTTCTTAAAGGCTGGGAGCCTGCTTTTTTGT |
| 14 | o-CENP-B _pE4XmnI | TCCACCATGGGAACCGTGTCCTGCAGCGGCGTGG |
| 15 | o-CENP-B_RV_pE4EcoRV | AAGCTGGGTCTAGATGCTAGCTTTGATGTCCAAGAC |
| 16 | o-HP1α_FW_pE4XmnI | TCCACCATGGGAACCATGGGAAAGAAAACCAAGCG |
| 17 | o-HP1α_RV_pE4EcoRV | AAGCTGGGTCTAGATGGTTTAAACGCTCTTTGCTG |
| 18 | o-HP1β_FW_pE4XmnI | TCCACCATGGGAACCATGGGGAAAAAACAAAAC |
| 19 | o-HP1β_RV_pE4EcoRV | AAGCTGGGTCTAGATGTTAGTTCTTGTCATCTTTTTTGTC |
| 20 | o-HP1β_N_upper | TGATTCTGAAATCTAGACCCAGCTTTCTT |
| 21 | o-HP1β_N_lower | TAGATTTCAGAATCAGAATCAGCTTTG |
| 22 | o-HP1β_C_upper | AGGCTCCGATAAGGGAGAGGAGAGCAAACC |
| 23 | o-HP1β_C_lower | CCCTTATCGGAGCCTGCTTTTTTGTAC |

| | |
|---|---|
| o-HP1γ_FW_pE4XmnI | TCCACCATGGGAACCATGGCCTCCAACAAAACTAC |
| o-HP1γ_RV_pE4EcoRV | AAGCTGGGTCTAGATGTTATTGAGCTTCATCTTCTGG |
| o-G9a/FW-pE4XmnI | TCCACCATGGGAACCATGGCGGCGGCGGCGGGAGCTGCAGCGGCGGCG |
| o-G9a/RV-pE4EcoRV | AAGCTGGGTCTAGATGTGTGTTGACAGGGGGCAGGG |
| o-G9a_775_FW | ACGAAAGGGGACCCCGGGTCCC |
| o-G9a_850_stop_RV | AGTGTCTGCTCTCCCTATTCAGACTTGCTGTCGGAGTC |
| o-G9a_889_FW | GCAGGCTCCACCATGGTTGAAGCTCTAACTGAAC |
| o-G9a_2542_RV | GGGGGTGTCCCCATGGTAGTT |
| o-AAVS1_MTdetect_FW | ATTGTCACTTTGCGCTGCCC |
| o-AAVS1_MTdetect_RV | AAGAGTGAGTTTGCCCAAGC |
| o-CENPB_MTdetect_FW | GATTCAGACAGTGAGGAAGAG |
| o-CENPB_MTdetect_RV | TGCTGTGGTTAGTCCACTGAG |
| o-HP1α_MTdetect_FW | TTTGAGACTCAAGAGCAGGG |
| o-HP1α_MTdetect_RV | AACGTAAGCTCCACAAGCGG |
| o-HP1β_MTdetect_FW | TTTGTGTGTGGCTACAGACC |
| o-HP1β_MTdetect_RV | CCCTTATATCTGTCCCCAG |
| o-HP1γ_MTdetect_FW | ATTTTGGTGGTGGGTTGTAAG |
| o-HP1γ_MTdetect_RV | TAGACCTCAAATGAGACACC |
| o-CENP-B_C_HR_R_FW | CATAGCTGTTTCCTGTCACTGGACCTAGCTGTGCC |
| o-CENP-B_C_HR_L_RV | TCCCGGGGAGCTCCATCCGCTTTGATGTCCAAGACCTC |
| o-Puro_3’_FW | CTGGTGCATGACCCGCAAGC |
| o-MK243_MTdetect_RV | GACCTTGATATGCTGCCTGC |

| | |
|---|---|
| o-mAID_5'_RV | TGGAGCTCCCCGGGATCCGGTG |
| M13_RV | GTCATAGCTGTTTCCTG |

**Supplemental Table S2.** Plasmids and guide RNA sequences designed for generating KO cells using the CRISPR-Cas9 system.

| Plasmid | Target gene | Sequence (5' → 3') |
|---|---|---|
| pTORA14AAVS1 | *AAVS1* | GGGGCCACUAGGGACAGGAU |
| pTORA14CENP-B_C | *CENPB* | GACAUCAAAGCUGAGUCACU |
| pTORA14HA4 | *HP1*$\alpha$ | AGCGGACAGCUGACAGUUCU |
| pTORA14HB7 | *HP1*$\beta$ | GCCCUCUGAUUUAUCUGUCU |
| pTORA14HG1 | *HP1*$\gamma$ | CGACAAAUUCUUCAGGCUCU |
